# Invasion genetics of the silver carp (*Hypophthalmichthys molitrix*) across North America: Differentiation of fronts, introgression, and eDNA detection

**DOI:** 10.1101/392704

**Authors:** Carol A. Stepien, Anna E. Elz, Matthew R. Snyder

## Abstract

The invasive silver carp *Hypophthalmichthys molitrix* escaped from southern U.S. aquaculture during the 1970s to spread throughout the Mississippi River basin and steadily moved northward, now reaching the threshold of the Laurentian Great Lakes. The silver carp is native to eastern Asia and is a large, prolific filter-feeder that decreases food availability for fisheries. The present study evaluates its population genetic variability and differentiation across the introduced range using 10 nuclear DNA microsatellite loci, sequences of two mitochondrial genes (cytochrome *b* and cytochrome *c* oxidase subunit 1), and a nuclear gene (ribosomal protein S7 gene intron 1). Populations are analyzed from two invasion fronts threatening the Great Lakes (the Illinois River outside Lake Michigan and the Wabash River, leading into the Maumee River and western Lake Erie), established areas in the southern and central Mississippi River, and a later Missouri River colonization. Results discern considerable genetic diversity and some significant population differentiation, with greater mtDNA haplotype diversity and unique microsatellite alleles characterizing the southern populations. Invasion fronts significantly differ, diverging from the southern Mississippi River population. About 3% of individuals contain a unique and very divergent mtDNA haplotype (primarily the southerly populations and the Wabash River), which may stem from historic introgression in Asia with female largescale silver carp *H. harmandi*. Nuclear microsatellites and S7 sequences of the introgressed individuals do not significantly differ from silver carp. MtDNA variation is used in a high-throughput sequence assay that identifies and distinguishes invasive carp species and their population haplotypes (including *H. molitrix* and *H. harmandi*) at all life stages, in application to environmental (e)DNA water and plankton samples. We discerned silver and bighead carp eDNA from four bait and pond stores in the Great Lakes watershed, indicating that release from retailers comprises another likely vector. Our findings provide key baseline population genetic data for understanding and tracing the invasion’s progression, facilitating detection, and evaluating future trajectory and adaptive success.

## Introduction

Discerning the population genetic trajectories of nonindigenous species invasions can enhance our overall understanding of the evolutionary adaptations and changing ecological community dynamics governing today’s ecosystems [1–3]. Establishments by invasive species comprise accidental experiments that ground-truth evolutionary and ecological theory with reality [4–6]. The genetic variation of invasive populations often affects their relative success and persistence in new habitats, including colonizing, reproducing, spreading, and overcoming biotic resistance [2, 7, 8].

### Invasion genetics

Invasion biology theory predicts that many introduced species would be characterized by low genetic diversity due to founder effect, which is believed to limit their adaptive potential (9–11). Countering this, large numbers of introduced propagules and multiple introduction events and sources may enhance the genetic diversity of invasions, with some actually possessing higher levels than found in native regions [12–14]. For example, genetic studies of North American Laurentian Great Lakes invasions resulting from ballast water introductions of the Eurasian zebra mussel *Dreissena polymorpha* (Pallas, 1771), quagga mussel *D. rostriformis* (Deshayes, 1838) [15, 16], and round goby *Neogobius melanostomus* (Pallas, 1814) [5, 17–19], all discerned relatively high genetic diversities, along with significant population differentiation across their introduced ranges. These results were attributed to large numbers of introduced propagules stemming from multiple introduction events, which traced to several Eurasian sources. Such cases of high genetic variability are believed to enhance the adaptive potential of invasions (see [11]).

Populations at the expansion fronts of an invasion front frequently possess less genetic variability than those at the core of the invasion, which is termed the “leading edge” hypothesis [20, 21]. The phenotypes of individuals at these front populations are postulated to be adapted for dispersal and high reproductive output [22], where they experience low population density, and may benefit from greater resource availability and less intraspecific competition, enhancing reproductive success [23, 24]. These processes may lead to genetic differences in expansion fronts versus longer-established populations, which may result from drift and/or selection. However, documentation of their comparative genetics across the spatial and temporal course of invasions are relatively rare in the literature [22].

Sometimes closely related invasive species, which may interact competitively, are introduced together or in close succession, as occurred in the Laurentian Great Lakes for the zebra and quagga mussels [25–27], with the latter overtaking the former in many areas [28, 29]. Another case of simultaneous introduction by related species include the Eurasian round and tubenose *Proterorhinus semilunaris* gobies [30], with the former continuing to be much more numerous and widespread in the Great Lakes [31]. In the 1960s and 1970s, the closely related east-Asian silver *Hypophthalmichthys molitrix* (Valenciennes 1884) and bighead *H. nobilis* (Richardson 1845) carps both were intentionally introduced to the southern U.S. to control algae [32]. The two species are known to hybridize [33, 34], which may increase their success via “hybrid vigor”. Silver and bighead carps were raised at six state, federal, and private facilities in Arkansas during the 1970s, and had been stocked into several municipal sewage lagoons [35]. They escaped in the 1970s and became established in the Mississippi River basin [32], spreading into the upper Mississippi River system [36]. The two species now are near the Great Lakes in the Illinois River system outside of Lake Michigan in the vicinity of Chicago IL, with growing concern that they will enter and become established in the Great Lakes [37, 38].

The Great Lakes already constitute one of the world’s most heavily invaded aquatic ecosystems, harboring over 186 reported invasive species, with the most significant ecological impacts caused by the invasions of the sea lamprey *Petromyzon marinus*, common carp *Cyprinus carpo*, zebra and quagga mussels, round goby, spiny waterflea, and rusty crayfish *Orconectes rusticus*. Ecologists have theorized that such a long history of established invasions is believed to increase the chances of success by subsequent invaders [39]. This may pave the way to success of the silver and bighead carps, with the grass carp *Ctenopharyngodon idella* now recently established in Lake Erie [40].

Accidental releases from aquaculture operations are a primary vector for introductions of exotic species [41], as led to the establishment of wild populations of silver and bighead carps in North America [32]. Other cases include the widespread escapes and establishments by tilapia (*Oreochromis spp*., *Sarotherodon spp*., and *Tilapia spp*.) [42], and cultured bivalves, including the Pacific oyster (*Crassostrea gigas*) in North America and Europe, and the Mediterranean mussel *Mytilus galloprovincialis* in South Africa [43].

### Invasion ecology of silver and bighead carps and study objectives

Silver and bighead carps are prolific filter-feeders that significantly alter and reduce planktonic community composition and availability for fisheries [44–47]. They often swim just below the surface, and can travel in large schools [48]. They mature between 4–8 years old in their native range, but as early as 2 years in North America, with females laying up to 5 million eggs per year [49]. The reproductive season appears to be more extensive in their invasive habitats in the Mississippi River system than in the native range [50]. The eggs are buoyant, hatch in a day, and the larvae readily disperse with currents [48]. Adults live to 20 years, reaching over 100 cm and 27.3 kg [48]. Modeling studies indicate that the introduction of even a small number of silver or bighead carps could establish successfully reproducing populations in the Great Lakes region, if individuals are of reproductive age and maturity [51].

The present geographic range for the silver carp invasion is illustrated in Fig. 1 [52]. Silver carp populations extend upstream and downstream of 23 locks and dams (three in the Arkansas River, seven in the Illinois River, eight in the Mississippi River, and five in the Ohio River) throughout the Mississippi River drainage. There currently are two potential man-made impediments to silver and bighead carps reaching the Great Lakes basin, the first being an electric barrier in the Chicago Area Waterway System separating the Illinois River from Lake Michigan, which is occasionally breached by small fishes and those caught in the wake of large boats [53]. In 2016, a 2.7km long, 2.3m -tall earthen berm was completed in Eagle Marsh, Fort Wayne, IN, a wetland that experienced flooding between the Wabash and Maumee Rivers, the latter leading into Lake Erie [54]. Populations at these two invasion front areas uniquely are analyzed in the present study. The issue of silver and bighead carps potentially entering and establishing in the Great Lakes has been of great concern to managers and the commercial and sport fishing communities [55, 56].

**Fig 1.**
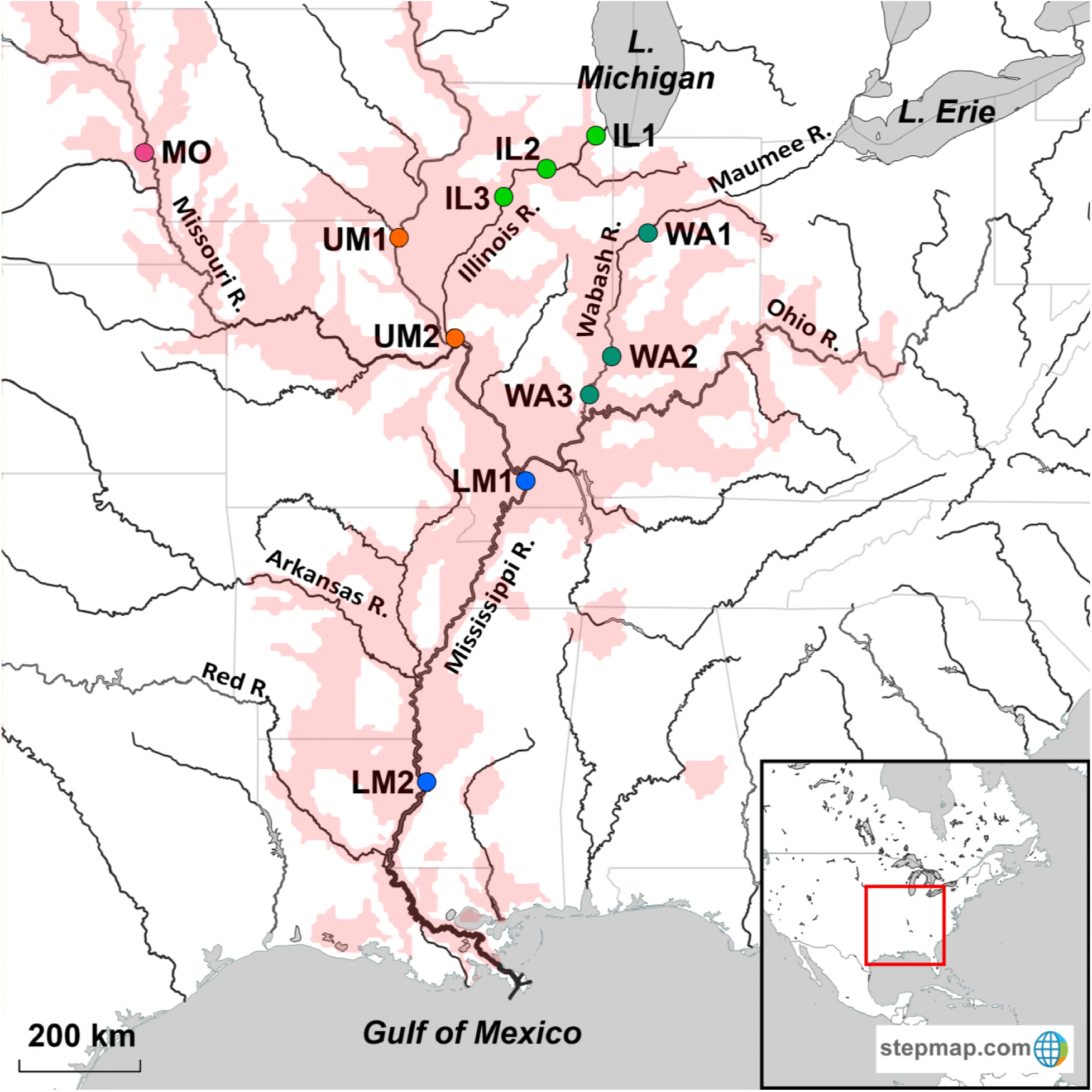
Map of silver carp invasive range (red) and sampling locations (circles, colored by waterway). (IL) Illinois R. IL1: LaGrange, IL (41.757623, −87.849964), IL2: Marseilles, IL (41.322269, −88.707172), IL3: Chillicothe, IL (40.929296, −89.461653). (WA) Wabash R. WA1: Lafayette, IN (40.430177, −86.898111), WA2: Vincennes, IN (38.688718, −87.526298), WA3: New Harmony, IN (38.135032, −87.940739). MO: Missouri R., Blair, NE (41.545091, −96.095555). (UM) Upper Mississippi. UM1: Warsaw, IL. (40.362244, −91.444024), UM2: Grafton, IL (38.966248, −90.430843). (LM) Lower Mississippi. LM1: Laketon, KY (36.868076, −89.124098), LM2: Vicksburg, MS (32.331320, −90.898628).

Currently, little is known about the population genetic patterns of the invasive silver and bighead carps in North America. The objective of the present study is to evaluate the genetic diversity and test for population structure of silver carp across the broad geographic range of its invasion, including the seat of original establishment in the Mississippi River basin and the two most likely invasion fronts leading to the Great Lakes, in the Illinois and Wabash Rivers. We test invasion genetics theory, including the possible presence of founder effect and correspondence to the leading edge hypothesis at invasion front areas. We further analyze whether genetic divergence and differentiation occurs across the invasive range, including at two primary invasion front areas. We evaluate diversity at 10 nuclear microsatellite (μsat) loci and sequenced two mitochondrial genes, for 922 base pairs (bp) of cytochrome *b* (cyt *b*) and ~600 bp of the cytochrome oxidase subunit I (COI). We compare those sequences with nuclear DNA sequencing variation from the single copy ribosomal protein S7 gene intron 1, and relate our results to other genetic studies of silver carp and other invasions.

## Materials and Methods

### Sample collection

Tissue samples from silver carp individuals were collected by our laboratory members and/or by federal agency, state agency, and university collaborators (see Acknowledgements) under state or federal collection permits, using agency protocols and the University of Toledo IACUC protocol #205400, “Genetic studies for fishery management” to CAS. Samples were collected from 11 population areas (Fig. 1, Table 1), representing much of the North American range of silver carp. Bighead carp also were sampled, along with grass carp, black carp *Mylopharyngodon piceus*, and common carp to provide comparative data from related species. Morphological characters, including gill-raker structure, were used to distinguish silver carp from bighead carp and discern possible hybrids (see [33]). Samples were labeled, stored in 95% EtOH, and archived at the NOAA Pacific Marine Environmental Laboratory.

**Table 1.** Sample locations and numbers of silver carp sequenced for each maker. *N* individuals processed for microsatellites (μsats) and DNA sequences of: cytochrome *b* (cyt*b*),COI, concatenated (concat) mitochondrial (both cytb and COI), and nuclear single-copy ribosomal protein S7 gene intron 1 (S7).

| **Population/Sample** | **Location** | $N_{\mu\text{sats}}$ | $N_{\text{cytb}}$ | $N_{\text{COI}}$ | $N_{\text{concat}}$ | $N_{\text{S7}}$ |
| --- | --- | --- | --- | --- | --- | --- |
| **Illinois River** | *Total* | 85 | 62 | 69 | 60 | 2 |
| IL1 | LaGrange, IL | 10 | 12 | 16 | 12 | 0 |
| IL2 | Marseilles, IL | 50 | 24 | 26 | 24 | 0 |
| IL3 | Chillicothe, IL | 25 | 26 | 27 | 24 | 2 |
| **Wabash River** | *Total* | 54 | 42 | 49 | 39 | 3 |
| WA1 | Lafayette, IN | 44 | 37 | 39 | 34 | 3 |
| WA2 | Vincennes, IN | 5 | 1 | 5 | 1 | 0 |
| WA3 | New Harmony, IN | 5 | 4 | 5 | 4 | 0 |
| **Missouri River** | Blair, NE | 50 | 29 | 30 | 29 | 3 |
| **Upper Mississippi River** | *Total* | 60 | 33 | 34 | 33 | 4 |
| UM1 | Warsaw, IL (Pool 20) | 10 | 9 | 10 | 9 | 2 |
| UM2 | <u>Grafton, IL (Pool 26)</u> | 50 | 24 | 24 | 24 | 2 |
| **Lower Mississippi River** | *Total* | 60 | 38 | 38 | 37 | 7 |
| LM1 | Laketon, KY | 10 | 10 | 10 | 10 | 1 |
| LM2 | Vicksburg, MI | 50 | 28 | 28 | 27 | 6 |
| *Total* |  | 309 | 204 | 220 | 198 | 19 |

### Population genetic data collection

Genomic DNA was extracted and purified from the EtOH fixed tissues using DNeasy^®^ Blood and Tissue Kits (Qiagen Inc., Valencia, CA USA), quality checked on 1% agarose mini–gels stained with ethidium bromide, and assessed with a Nanodrop^™^ spectrophotometer (Thermo Scientific, Bothell, WA, USA). Genetic variation was analyzed at 10 nuclear DNA microsatellite loci, including *Hmo1* and *Hmo11* from Mia *et al.* 2005 [57], *Ar201* from Cheng *et al.* 2007 [58], and *HmoB4, B5, D8, D213, D240, D243*, and *D246* from King et al. 2011 [59]. Representative subsets, which included individuals from each location and microsatellite variants, then were sequenced for the mitochondrial (mt)DNA cytochrome oxidase subunit I (COI) and cytochrome *b* (cyt*b*) genes and for the nuclear ribosomal protein S7 gene intron 1 (S7).

For the microsatellites, 10μL polymerase chain reactions (PCR) contained 0.35 units AmpliTaq^®^ DNA polymerase (ABI; Applied Biosystems^™^, Foster City, CA, USA), 1X GeneAmp^®^ PCR buffer I (ABI), 80μM total dNTPs, 0.4mM spermadine, 0.52μM of each primer, and 2μL of ≥30ng/μl DNA and ddH2O. PCR was conducted on C1000^™^ thermal cyclers (Bio–Rad Laboratories, Hercules, CA) with 2 min initial denaturation at 94°C, followed by 39 cycles of 40 sec denaturation at 94°C, 40 sec annealing (at 52°C for the Mia *et al.* 2005 primers [57], 58°C for the Cheng *et al.* 2007 primers [58], and 56°C for the King *et al.* 2011 [59] sets), and 1 min 72°C extension, capped by 10 min final 72°C extension. Amplification products were diluted 1:50 with ddH2O, of which 2μL was added to 13 μL solution of formamide and ABI GeneScan^™^-500 LIZ^®^ size standard, and loaded into 96-well plates. Microsatellite products were denatured for 2 min at 95°C and analyzed on our ABI 3130xl Genetic Analyzer with GeneMapper^®^ 4.0 software (ABI). Output profiles were checked manually to confirm allelic size variants.

We aligned and analyzed 992 bp of sequence for the mtDNA cytb gene, which was amplified using the primers Song-F (5′–GTGACTTGAAAAACCACCGTTG–3′) [60] and H5 (5′–GAATTYTRGCTTTGG-GAG–3′) [61] and 549 bp of the COI gene, using COIFF2d (5′–TTCTCCACCAACCACAARGAYATYGG–3′) and COIFR1D (5′–CACCTCAGGGTGTC CGAARAAYCARAA–3′) [62]. PCR reactions contained 25μL of 1.25 units AmpliTaq^®^ DNA polymerase, 1X GeneAmp^®^ PCR Buffer I, 250μM dNTPs, 0.5uM (cyt*b*) or 1μM (COI) of each primer, and 2μL of ≥30ng/μL of DNA template, and ddH_2_O. Conditions were 3 min at 94°C, followed by 34 cycles of 95°C for 30 sec, 40 sec at 50°C, and 72°C for 45 sec, capped by 5 min at 72°C. We also sequenced the S7 intron using S7RPEX1F and S7RPEX2R following Chow and Hazama 1998 [63] for a representative subset of silver carp individuals, which included all mtDNA haplotypes (see Results), representatives of all populations, bighead carp, and other carp species.

PCR product aliquots (4μL) were visualized on 1% agarose mini-gels stained with ethidium bromide and successful reactions were purified with QIAquick^®^ PCR Purification Kits (Qiagen) and quantified via Nanodrop. Sanger DNA sequencing was outsourced to the Cornell University Life Sciences Core Laboratories Center (http://cores.lifesciences.cornell.edu/brcinfo/) and MCLAB (http://www.mclab.com/DNA-Sequencing-Services.html), which used ABI Automated 3730 DNA Analyzers. Sequences were quality scored, manually checked, and aligned by us with Codon Code Aligner v7.01 (Codon Corp.). Additional DNA sequences were mined from N.I.H. GenBank (provided in Table S1).

### eDNA assay development, sampling and testing

Using the mtDNA sequence variation we discerned for the cyt*b* gene, we developed two high-throughput Illumina^®^ MiSeq assays to facilitate detection and quick differentiation of silver and bighead carps and other related cyprinid species in field samples, following our published laboratory procedures [64]. Assay one used forward primer 5’–CACACNTCNAAACAACGAG GNCTNACNTTCCG–3’ and reverse primer 5’–GGGTGTTCNACNGGYATNCCNCCAAT TCA–3’ to amplify a 55 base pair (bp) region from cytb nucleotides 954–1008. This assay distinguishes among all invasive carp species, as well as the highly divergent silver carp haplotype “H” (see Results), and two other groups of silver type haplotypes (“A–E” from “F” and “G”). Our assay two used forward primer 5’–AACBCARCCBCTBCTWAAAATRGC–3’ and reverse primer 5’–AYTCANCCYAAYTTWACYTCWCGGC–3’ to amplify a 167 bp region spanning nucleotides 42–208, which identifies and discerns among all invasive carp species. It also distinguishes among five of the silver carp haplotype variants, differentiating the “B”, “E”, and “H” haplotypes, grouping together haplotypes “A”, “C”, and “D”, and grouping together “F” and “G”. Illumina^®^, MiSeq sequencing adapters, indices, and P5/P7 adapters were added to the ends of amplicons. The 25μl PCR reactions consisted of 1x Radiant TAQ Reaction Buffer (Alkali Scientific Inc., Ft. Lauderdale, FL), 3mM MgCl2, 1mM total dNTPs, 0.6mM of each primer (with an Illumina^®^, MiSeq sequencing primer tail), and 1.25 units of Radiant TAQ polymerase. The final PCR step used the prior step’s column-cleaned product as template (2μl) and incorporated Nextera paired end indices (Illumina^®^, kit FC-121–1011), which included the P5 and P7 adaptor sequences, allowing the prepared library to bind onto the surface of the Illumina^®^ MiSeq flowcell. Adding unique index combinations permitted multiple samples to be pooled together in each MiSeq lane. No template controls (NTCs) were amplified for each assay, and only libraries that lacked NTC amplification were used for MiSeq analyses (see below).

After column clean up, products were sized and quantified in our laboratory on a 2100 Bioanalyzer (Agilent Technologies), prior to Illumina^®^ MiSeq analysis by Ohio State University’s Molecular and Cellular Imaging Center in Wooster, OH (http://mcic.osu.edu/). To avoid sequencing dimer product observed around 200–250 bp, the targeted fragments (~350 bp) were size-selected with a 1.5% agarose gel cassette on Pippin Prep (Sage Science). Concentrations of pooled products were measured with a Qubit fluorometer (Invitrogen). Pooled samples were run on an Illumina^®^ MiSeq with 2X 300 bp V3 chemistry. An additional 40–50% PhiX DNA spike-in control was added to improve data quality of low nucleotide diversity samples, per our published laboratory protocol [64].

We then used these assays to test for the possible cryptic presence of silver and bighead carps and other related cyprinid species in retail bait shops and pond supply stores, as part of a larger ongoing project. We evaluated water samples and live bait from 45 bait shops and 21 pond stores located in the Lake Erie, Lake St. Clair, and Wabash River watersheds in 2016 and 2017. Approximately two dozen bait fish were purchased from each bait shop, which were immediately sacrificed in the parking lot using the University of Toledo IACUC procedure. The water containing the fish was drained into a sterile container, placed on ice, and then frozen at −80°C in the laboratory. We also sampled water from the pond stores, which variously sold plants, snails, and fishes (primarily koi). We centrifuged 250 ml of the water from each shop at 4500 rpm and 4°C, for 45 min. The supernatant was poured off and the pellet, containing extra and intracellular DNA and debris, was resuspended in 95% ethanol and stored at −20°C until DNA extraction using Qiagen DNeasy kits. All bait and pond store sample extractions were amplified with the two assays. Species detections were considered valid if they either occurred above the error rate calculated from positive controls (see below) or if a species was discerned in both assays at any proportion.

### Microsatellite analyses

All loci were evaluated for linkage disequilibrium and conformance to Hardy-Weinberg equilibrium expectations, using the Markov Chain Monte Carlo (MCMC) procedure with 10,000 dememorizations, 1,000 batches, and 10,000 iterations per batch in GENEPOP v4.0 [65]. Significance values were adjusted with standard Bonferroni correction [66]. MICRO-CHECKER v2.2.3 [67] was used to examine results for possible scoring errors, large allele dropout, stuttering, and/or null alleles at each locus.

*F*STAT v. 2.9.3.2 [68] was employed to calculate measures of genetic diversity, including number of alleles per locus (*N*_A_), observed heterozygosity (*H*_O_), and allelic richness (*A*_R_), which were adjusted for sample size using rarefaction. Standard errors were calculated separately in EXCEL (Microsoft, Redmond, WA) and are presented as ±S.E. Significance values (*p*<*a*) were adjusted with sequential Bonferroni correction [69]. *H*_O_ and *A*_R_ were evaluated for significant differences with Friedman rank sum tests in R v.3.2.1 [70]. Number of private alleles (*N_PA_*) per locus, i.e., those appearing unique to a population sample were identified using CONVERT v1.31 [71]. Percentage of private alleles (*P_PA_*) was determined by dividing the number of private alleles for a given sample by its total number of alleles. Due to disparity in sample size, the rarefaction representation of private alleles was evaluated with the program ADZEv1.0 [72]. COLONY v.2.0.6.1 [73] was used to test for the possible presence of siblings in the dataset. Evidence for selection was evaluated using the outlier method of Beaumont and Nichols 1996 [74] in LOSITAN [75].

Pairwise genetic divergences were calculated between all population samples, and separately between the two invasion fronts at the Illinois and Wabash Rivers using the *F*_ST_ analog *θ*_ST_ (76) in *F*STAT, which is regarded as appropriate for analyzing high gene flow species, small sample sizes, and unknown number of subpopulations [77–79], and to facilitate comparisons with other studies. Since *F*-statistic estimates assume a normally distributed data set [76] and may be influenced by sample size [80], we additionally conducted pairwise exact tests of differentiation (χ^2^) in GENEPOP, using MCMC chains of 10,000, 1000 batches, and 10,000 iterations. Probability values for both types of pairwise comparisons were adjusted using sequential Bonferroni correction [69]. This correction is regarded as a very conservative approach that may preclude elucidation of significance when sample sizes are low, leading to type II error (i.e., falsely rejecting the null hypothesis of no significant difference between samples [81]. Thus, we report significance values both after (**) as well as prior to (*) sequential Bonferroni correction, so that results on the borderline can be discerned (which may have been influenced by sample size limitations).

In addition, *θ*_ST_ and exact tests of differentiation also were conducted to compare all individuals possessing highly divergent mtDNA haplotype “H” (see Results) with 14 other individuals, which included all other haplotypes (e.g., “A–G”). The purpose was to determine whether the nuclear DNA of those individuals also was genetically divergent using microsatellites. Those individuals also were sequenced for the nuclear S7 gene intron.

Population relationships from microsatellite variation were visualized using three-dimensional factorial correspondence analysis (3D-FCA) [82] in GENETIX v4.05 [83]. Bayseian STRUCTURE v. 2.3.2 [84] analysis was used to determine whether, and how many, discrete genetic groups were represented. Ten replicates evaluated scenarios from *K*=1–6, with burn ins of 50,000 and 100,000 iterations, and relative support for each was determined using delta *K* [85] in STRUCTURE HARVESTER [86]. Individual assignment tests were conducted with GENECLASS2 [87] for all populations sampled, and separately for the invasion fronts at the Illinois and Wabash Rivers, using the “enable probability computation” option, the simulation algorithm from Paetkau *et al.* 2004 [88], and 100,000 simulations.

### DNA sequence analyses

We evaluated mtDNA sequence variation in the cyt*b* and COI regions, both separately and together as concatenated sequences (Table 1). We incorporated complete sequences for the gene regions from NIH GenBank (https://www.ncbi.nlm.nih.gov/genbank/; detailed in Supplementary Table 1). Our sequences are deposited in GenBank as Accession numbers XXXX–XXXX. Haplotype relationships were depicted with TCS haploptype networks [89] constructed in POPART (http://popart.otago.ac.nz), in reference to four bighead carp haplotypes. ARLEQUIN was used to calculate the numbers of haplotypes (*N*_H_), haplotypic diversity (*h*), numbers and proportions of private haplotypes (*N*_PH_ and *P*_PH_) per sample, and pairwise divergences between population samples using *θ*_ST_ and exact tests [90]. Relative percentage of haplotypes per population sample was illustrated using a stacked bar graph, drawn with R. Bayesian phylogenetic trees were used to evaluate the relationships among the mtDNA concatenated and the nuclear S7 sequence haplotypes in MRBAYES v.3.2.6 [91], in comparison with representative sequences from bighead carp, grass carp, black carp, and common carp. We used the GTR + I + Γ model of substitution, a relaxed molecular clock, and the rate at which the variance of the effective branch length increases over time, using the default prior exponential distribution rate =10.0. Common carp was the selected outgroup. The MCMC was run for 100,000 generations to calculate support values for the branch nodes. FIGTREE v.1.4.3 (http://tree.bio.ed.ac.uk/software/figtree/) was used to visualize the consensus tree.

### High-throughput sequencing data analyses

Custom bioinformatic scripts in PERL v5.26.1 were employed to trim the primers from the FASTQ files returned from the Illumina^®^, MiSeq runs. Trimmed sequences were merged, dereplicated (grouped by 100% sequence similarity), and chimeras were removed using the program DADA2 [92]. DADA2 employed a denoising algorithm in a Poisson model to calculate the error probabilities at each nucleotide position. The algorithm removed variants having nucleotide changes that occurred at a lower percent representation than was predicted from their associated probabilities. Positive controls were run for each assay using samples prepared with known sequences. In those samples, unexpected operational taxonomic units (OTUs) that remained after the DADA2 de-noising algorithm was applied, were attributed to PCR and/or sequencing error or to sample mis-assignment (via index hopping). The greatest percent representation of any single erroneous OTU was set as the cutoff. The remaining OTUs were compared using the Basic Local Alignment Search Tool (BLAST; https://www.ncbi.nlm.nih.gov/) with the command line to a custom database containing cytb sequences of all fishes native to the Great Lakes basin [93] and non-native species that are present or are predicted future invaders [94]. All BLAST results with the lowest e value (best match) per OTU were summarized with a custom PERL script that grouped together identical or closely related reads from species. All μsat allele, Sanger, and high-throughput sequencing data were deposited in the DRYAD public data base (doi.org/XXXXX/dryad.XXX).

## Results

### Microsatellite diversity, divergence, and population structure

Results showed no linkage disequilibrium, null alleles, or selection. A single locus (*Hmo-B4*) that did not conform to Hardy-Weinberg equilibrium expectations was removed from further analyses. There were no differences in genetic diversity and *θ*_ST_ values when population samples were pooled regionally (e.g., all samples from the Lower Mississippi together, *N*=60) versus separately for individual sampling locations (Table 2; Fig. 1). Thus, results are presented with regional pooling to increase sample size and better represent overall geographic patterns across the range. Four pairs of full siblings were identified from the COLONY analyses for silver carp, whose inclusion or exclusion did not significantly alter *θ*_ST_; thus, the complete data are presented. One individual from three of the pairs was collected in the Illinois River, and both individuals of the fourth pair occurred in the Illinois River. Of the three located in separate populations, the other sibling was found in the Lower Mississippi River, the Upper Mississippi River, and the Missouri River, respectively. This indicates high dispersal of full siblings.

**Table 2.**
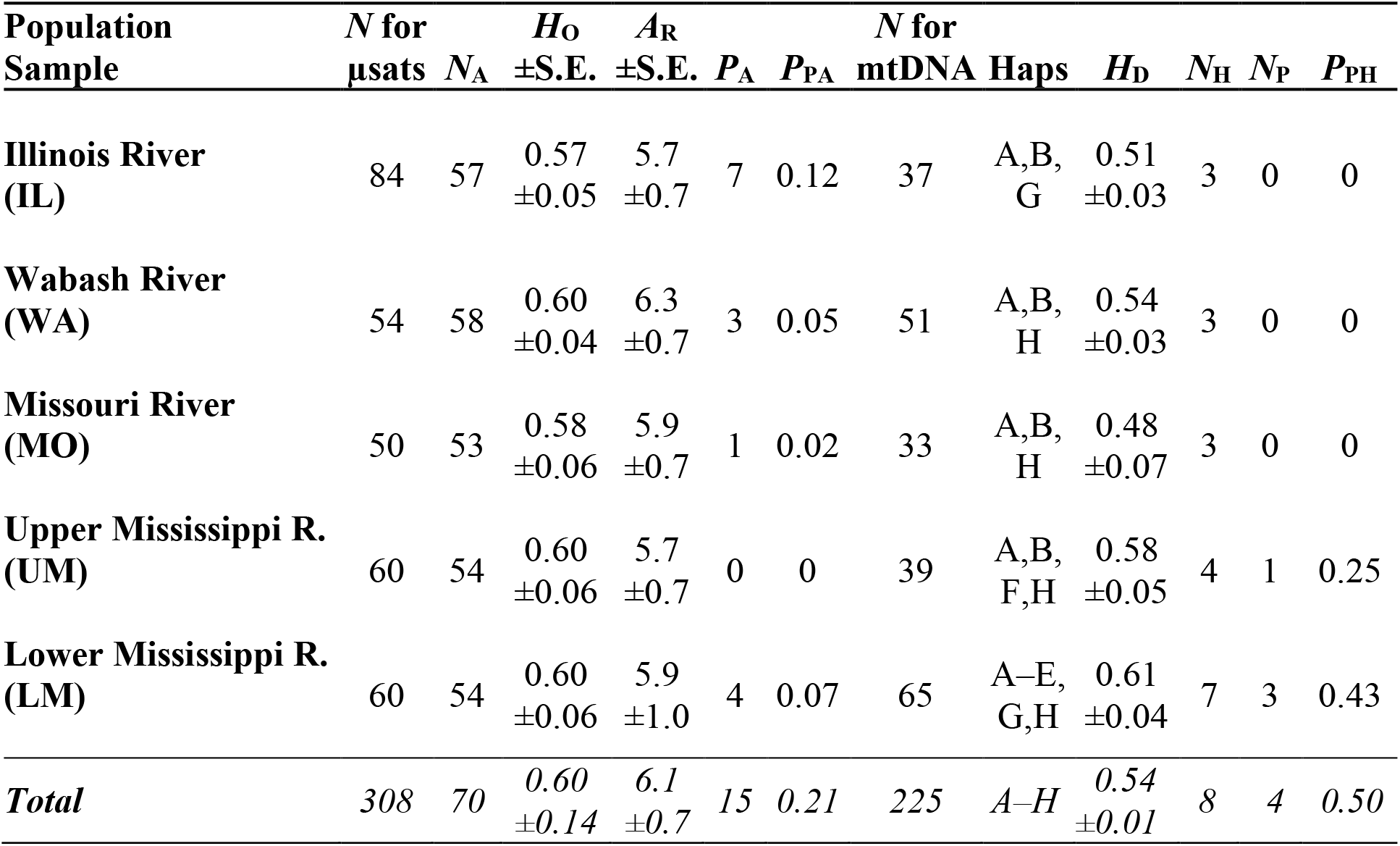
Genetic diversity of silver carp population samples. Number of individuals processed for microsatellites (*N* μsats) and mtDNA (*N* mtDNA), number of alleles/concatenated haplotypes (*N*_A_/*N*_H_), observed heterozygosity (*H*_O_) plus or minus standard error (±S.E.), and allelic richness (*A*_R_) ±S.E., number of private alleles/private haplotypes (*N*_PA_/*N*_PH_), proportion of alleles/haplotypes that are private to the sample (*P*_PA_/*P*_PH_), number of haplotypes (*N* Haps), and haplotypic diversity (*H*_D_).

| **Population Sample** | ***N* for μsats** | ***N<sub>A</sub>*** | ***H<sub>O</sub>* ±S.E.** | ***A<sub>R</sub>* ±S.E.** | ***P<sub>A</sub>*** | ***P<sub>PA</sub>*** | ***N* for mtDNA** | **Haps** | ***H<sub>D</sub>*** | ***N<sub>H</sub>*** | ***N<sub>P</sub>*** | ***P<sub>PH</sub>*** |
| --- | --- | --- | --- | --- | --- | --- | --- | --- | --- | --- | --- | --- |
| **Illinois River (IL)** | 84 | 57 | 0.57 ±0.05 | 5.7 ±0.7 | 7 | 0.12 | 37 | A,B,G | 0.51 ±0.03 | 3 | 0 | 0 |
| **Wabash River (WA)** | 54 | 58 | 0.60 ±0.04 | 6.3 ±0.7 | 3 | 0.05 | 51 | A,B,H | 0.54 ±0.03 | 3 | 0 | 0 |
| **Missouri River (MO)** | 50 | 53 | 0.58 ±0.06 | 5.9 ±0.7 | 1 | 0.02 | 33 | A,B,H | 0.48 ±0.07 | 3 | 0 | 0 |
| **Upper Mississippi R. (UM)** | 60 | 54 | 0.60 ±0.06 | 5.7 ±0.7 | 0 | 0 | 39 | A,B,F,H | 0.58 ±0.05 | 4 | 1 | 0.25 |
| **Lower Mississippi R. (LM)** | 60 | 54 | 0.60 ±0.06 | 5.9 ±1.0 | 4 | 0.07 | 65 | A–E,G,H | 0.61 ±0.04 | 7 | 3 | 0.43 |
| **Total** | 308 | 70 | 0.60 ±0.14 | 6.1 ±0.7 | 15 | 0.21 | 225 | A–H | 0.54 ±0.01 | 8 | 4 | 0.50 |

A total of 70 microsatellite alleles were discerned for silver carp (Table 2), with the greatest numbers at the invasion fronts, including 57 in the Illinois River and 58 in the Wabash River, versus 54 for the Lower Mississippi River population. Observed heterozygosity was 0.60 across all population samples, ranging from 0.57 in the Illinois River, to 0.60 in the Wabash River and the Upper and Lower Mississippi Rivers. Heterozygosity values did not significantly differ among the populations. Allelic richness appeared highest in the Wabash River front population at 6.3 and next highest in the Lower Mississippi River at 5.9, ranging down to 5.7 in the Illinois and Upper Mississippi River populations. No significant differences in allelic richness occurred among the populations. A total of 15 private alleles were identified, constituting 21% of the overall alleles (Table 2). Of these, the Illinois River invasion front possessed the most (seven, totaling 12% of its alleles), followed by the Lower Mississippi River (four at 7%), and the Wabash River (three at 5%).

Significant genetic divergences distinguished between all but two pairs of populations, assessed with the exact tests (Table 3). The two exceptions were similarities between the Upper Mississippi River versus the Missouri and Illinois river populations. Analyses using *θ*_ST_ divergences revealed two significant differences after sequential Bonferroni correction, between the Lower Mississippi River population versus the Missouri and the Illinois Rivers (*F*_ST_=0.012 and 0.007, respectively). All other comparisons were on the borderline (were significant prior to correction, but not after). Divergence between the two invasion front populations, Illinois and Wabash Rivers, was significant in all types of analyses (*F*_ST_=0.009, *p*<0.02, and exact test *p*<0.001; Table 3C).

**Table 3.** **Pairwise divergences, based on nuclear DNA microsatellite data, with θ_ST_** (below diagonal) **and exact tests** (above) **for: (A) all populations sampled** and **(B) the invasion front populations alone**. NS=not significant; *=*p*<0.05; **=remained significant after sequential Bonferroni correction.

| **A** |  |  |  |  |  |
| --- | --- | --- | --- | --- | --- |
| **Population Sampled** | **IL** | **WA** | **MO** | **UM** | **LM** |
| **Illinois R. (IL)** | ~ | ** | ** | NS | ** |
| **Wabash R. (WA)** | 0.009* | ~ | ** | ** | ** |
| **Missouri R. (MO)** | 0.005* | 0.008* | ~ | NS | ** |
| **Upper Mississippi R. (UM)** | 0.001* | 0.010** | 0.001* | ~ | ** |
| **Lower Mississippi R. (LM)** | 0.007** | 0.009** | 0.012** | 0.006* | ~ |
| **B** |  |  |  |  |  |
| **Location Sampled** | **IL** | **WA** |  |  |  |
| **Illinois R. (IL)** | ~ | ** |  |  |  |
| **Wabash R. (WA)** | 0.009** | ~ |  |  |  |

The 3-d FCA (Fig. 2) explained 88.54% of the data, distinguishing among all of the populations (Table 3). All populations appeared widely separated, with the greatest distinctions appearing for the Upper Mississippi, Illinois, and Wabash Rivers. STRUCTURE and STRUCTURE HARVESTER yielded the greatest support for *K*=2 population groups (not shown), for which distribution plots were uninformative (i.e., all individuals and samples showed mixed assignment to both hypothetical genetic groups). GENECLASS2 results for the entire data set also discerned overall mixed assignments. However, the GENECLASS2 assignment test between the two invasion front populations, the Illinois versus the Wabash rivers, indicated that 62% and 98% of their individuals respectively self-assigned. Individuals from the Wabash River population thus showed very high self-assignment.

**Fig 2.**
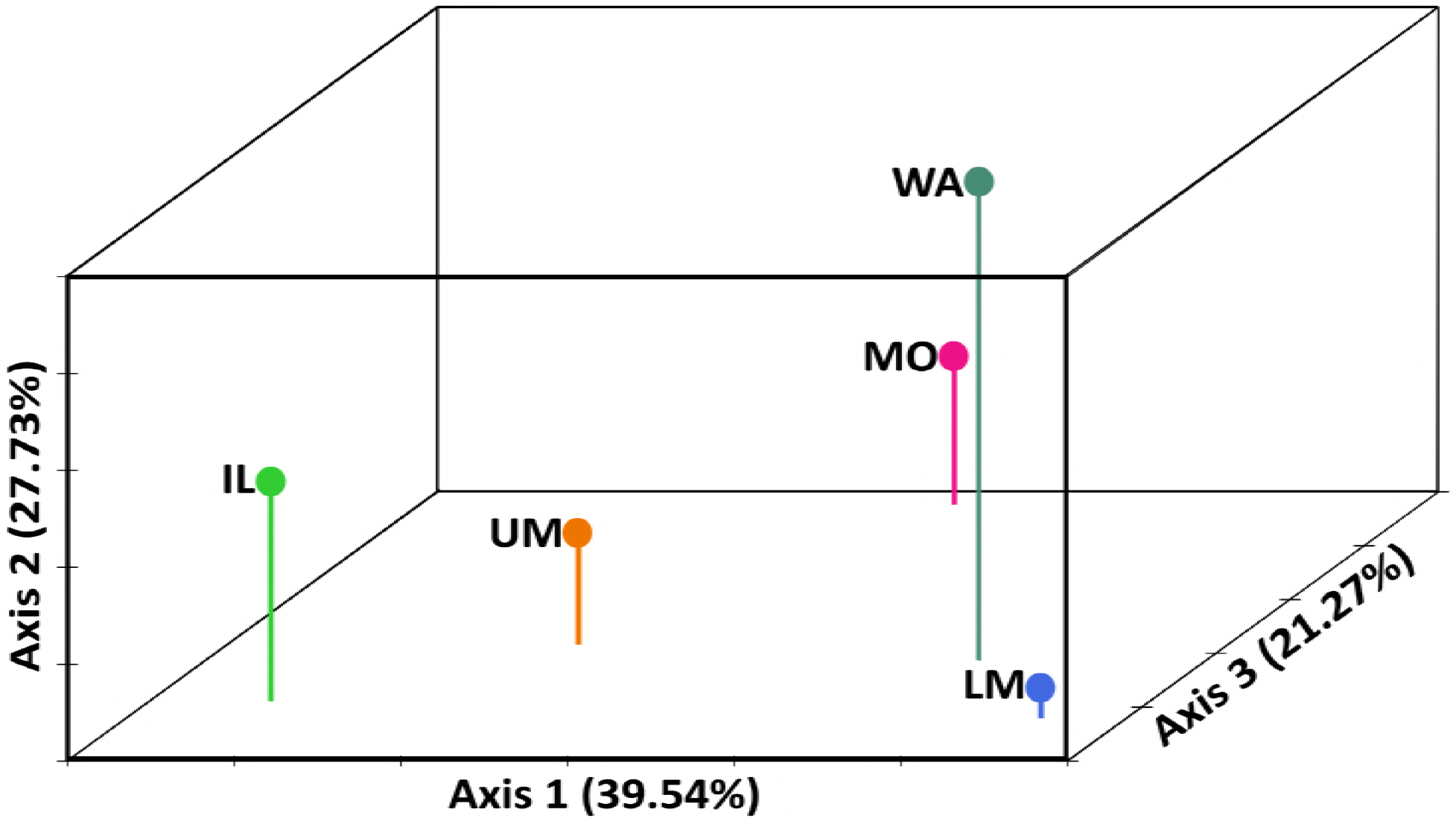
Three-Dimensional Factorial Correspondence analysis illustrating the genetic relationships among silver carp populations, based on the microsatellite results.

### Mitochondrial and nuclear DNA sequence haplotypes

In total, we analyzed 238 cyt*b* sequences (including 252 COI, and 230 concatenated mtDNA COI and cyt*b* sequences) for silver carp (Table 1). Accession numbers for sequences mined from GenBank are shown in the Supplementary Table S1. The cyt*b* dataset contained eight silver carp haplotypes in North America, which are lettered A–G (Fig. 3). Overall, 10 silver carp cytb haplotypes are known, which included two others from GenBank (Accession #AB198974, lettered “R” from the Black River in Russia and #AF051866, lettered “Q” from the Yangtze River in China). Their genetic relationships to the other cyt*b* haplotypes are depicted in Fig. 4A. Those two haplotypes were not included in our COI or concatenated datasets, which then numbered four and eight haplotypes respectively (Fig. 4B). New and unique haplotypes discerned in our study were deposited in GenBank (cytb:XXXX–XXXX and COI:XXXX–XXXX, S1 Table and Table 4). The remaining results are focused on the concatenated dataset.

**Fig 3.**
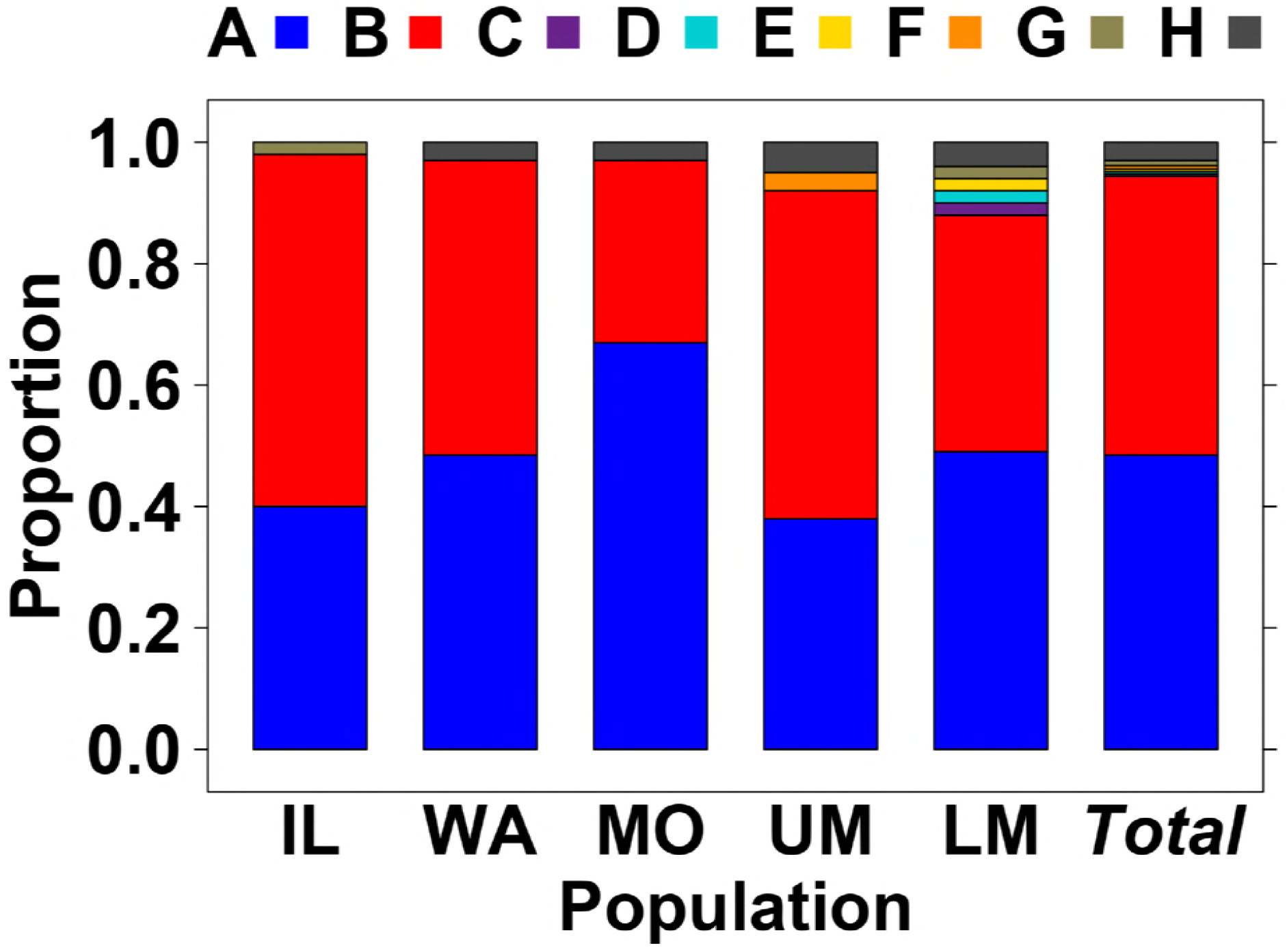
Silver carp mtDNA cytochrome *b* haplotype frequencies in North American populations. IL=Illinois River, WA=Wabash River, MO=Missouri River, UM=Upper Mississippi River, LM=Lower Mississippi River.

**Fig 4.**
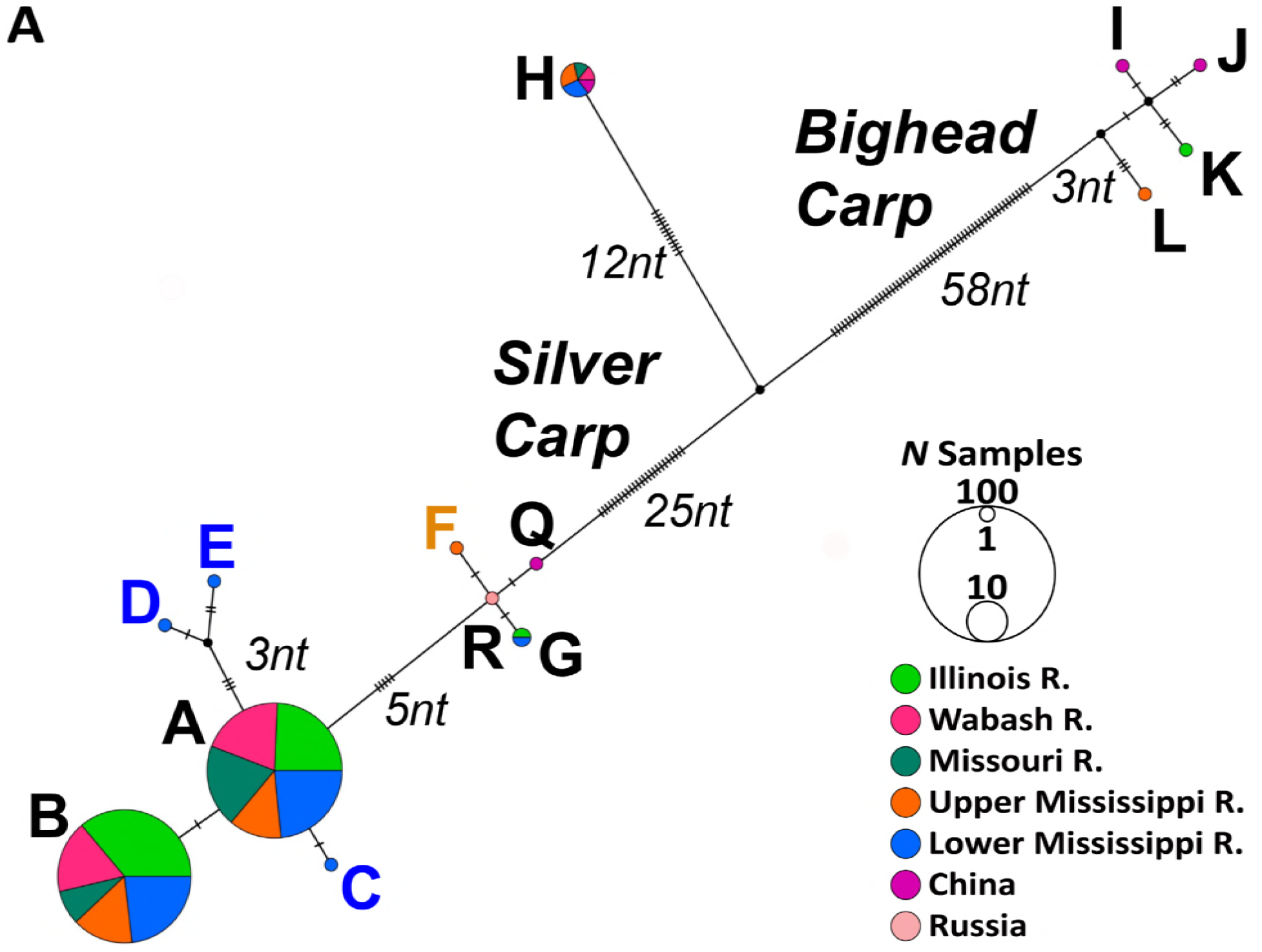
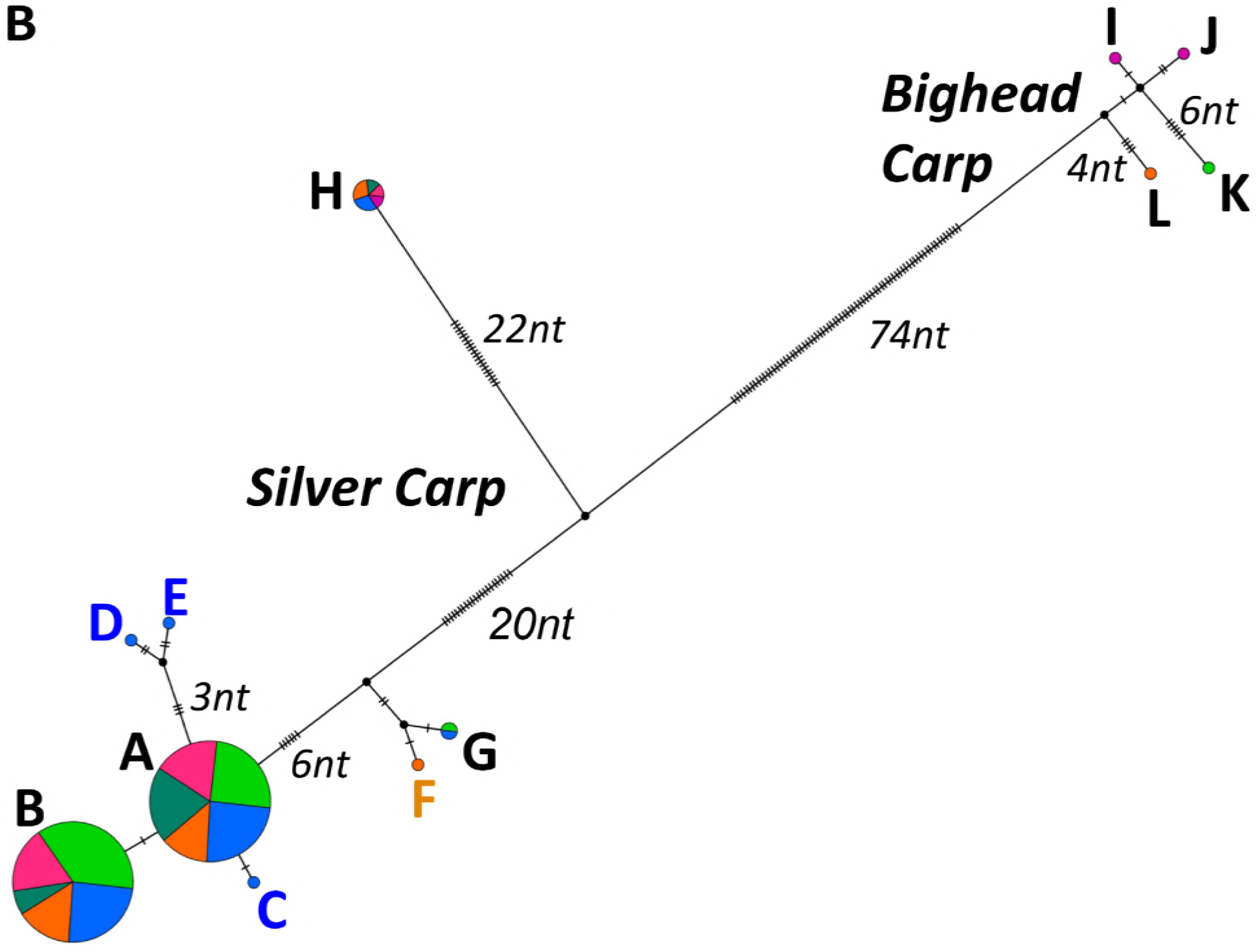
Silver carp TCS mtDNA haplotype networks of (A) cytochrome (cyt) *b* and (B) cyt *b* and COI concatenated mitochondrial DNA sequences. The (A) network for cyt-*b* contains GenBank sequences from Russia and China (haplotypes “R” and “Q”, respectively), which are not included in the concatenated network (B) because their COI sequences were unavailable. Bighead carp sequences (“I-L”) are included for comparison.

**Table 4.**
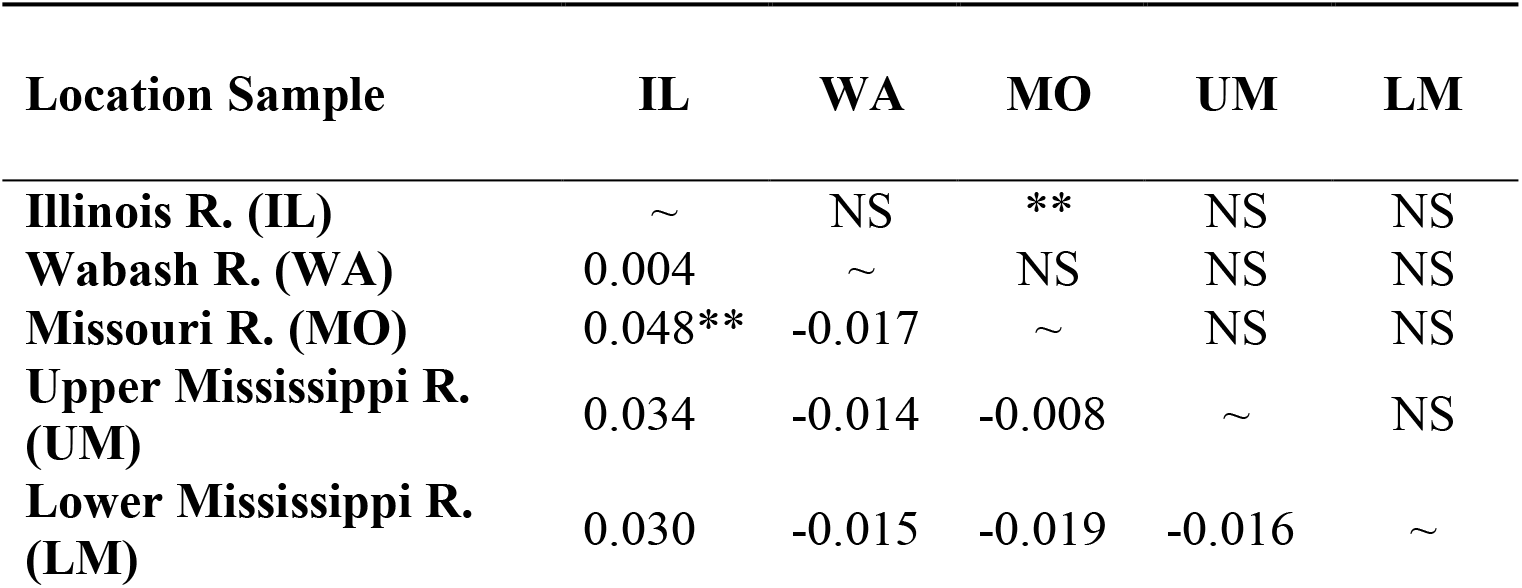
Pairwise divergences using concatenated mtDNA sequences. *θ*_ST_ (below diagonal), exact tests (above). *=*p*<0.05. **=significant after sequential Bonferroni correction.

| Location Sample | IL | WA | MO | UM | LM |
| --- | --- | --- | --- | --- | --- |
| **Illinois R. (IL)** | ~ | NS | ** | NS | NS |
| **Wabash R. (WA)** | 0.004 | ~ | NS | NS | NS |
| **Missouri R. (MO)** | 0.048** | -0.017 | ~ | NS | NS |
| **Upper Mississippi R. (UM)** | 0.034 | -0.014 | -0.008 | ~ | NS |
| **Lower Mississippi R. (LM)** | 0.030 | -0.015 | -0.019 | -0.016 | ~ |

Eight silver carp mtDNA concatenated haplotypes were identified in our North American samples (Fig. 3, Tables 2 and 4–5), with the most (seven) occurring in the Lower Mississippi River population (only “F” was not found in that population). The samples were overwhelmingly dominated by the common haplotypes “A” and “B” (Fig. 3), which together comprised over 90% of the samples. Haplotype “B” was less abundant at the Missouri River location than in the other populations. All populations contained haplotypes “A” and “B”, and all but the Illinois River also contained rarer haplotype “H”. Haplotypes “C”, “D”, and “E” were found only in the Lower Mississippi River population. Haplotypes “C”, “D”, and “E” were singletons, which evolutionary analysis depicted as most closely related to common haplotype “A” (Figs. 4 and 5A). Haplotypes “D” and “E” shared a closer relationship with each other (Fig. 5A). Haplotype “F” (a single individual) was discerned in the Upper Mississippi River population alone (Figs. 3–4). The invasion front populations of the Illinois and Wabash Rivers contained just three haplotypes each, with the Illinois River containing “G” and the Wabash River with “H”, and the former containing a greater abundance of haploytpe “B”. The sole differentiation at the population level in mtDNA pairwise divergence tests was between the Missouri River and the Illinois River samples (Table 4).

**Fig 5.**
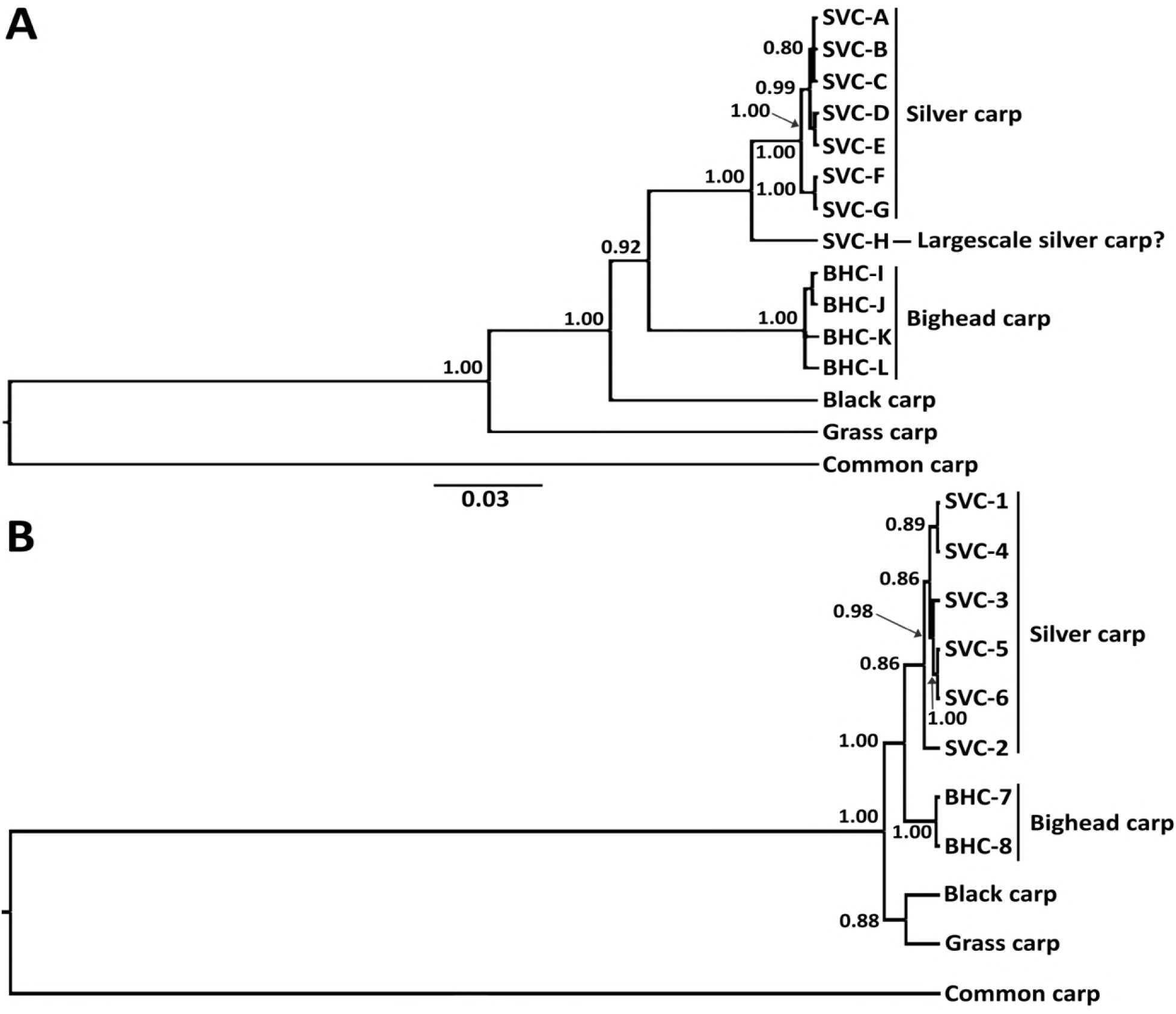
Bayesian trees of silver carp (SVC) and bighead carp (BHC) sequence haplotypes, in relation to black, grass, and common carp. (A) Concatenated mitochondrial DNA (mtDNA) tree. (B) S7 nuclear DNA tree. Incongruence with mtDNA tree implies a historical mtDNA introgression, possibly with the closely related largescale silver carp. SVC S7-“1” contained 12 individuals with haplotypes “A-H” (two having haplotype “A”, one individual each with the “B, C, D, E, F”, and “G” haplotypes, and four having haplotype “H”), S7-“2” (one individual with haplotype “A”), S7-“3” (one individual with haplotype “A”), S7-“4” (three individuals with haplotype “B”), S7-“5” (one individual with haplotype “G”), and S7-“6” (one individual with haplotype “H”).

**Table 5.** MtDNA concatenated haplotype (Hap) designations for sequence position variations of silver carp in the cytochrome *b* and COI gene regions. Based on nucleotide positions (above sequence) within the complete cytochrome *b* (bp 76–1136 1025), and COI (bp 66–447; in italics) gene regions. Dots indicate base identical to haplotype “A”. Only variable bases are shown. Haplotype “H” = putative *H. harmandi* introgression (all individuals were identical).

| Hap | 76 | 92 | 101 | 107 | 137 | 139 | 152 | 197 | 241 | 278 | 284 | 302 | 305 | 311 | 335 | 344 | 362 | 397 | 398 |
| --- | --- | --- | --- | --- | --- | --- | --- | --- | --- | --- | --- | --- | --- | --- | --- | --- | --- | --- | --- |
| A | C | G | G | A | C | A | T | T | G | C | G | A | G | A | T | C | G | G | A |
| B | . | . | . | . | . | . | . | . | . | . | . | . | . | . | . | . | . | . | . |
| C | . | . | . | . | . | . | . | . | . | T | . | . | . | . | . | . | . | . | . |
| D | . | . | . | . | . | . | . | . | A | . | . | . | . | . | . | . | . | . | G |
| E | . | . | . | . | . | . | . | . | A | . | . | . | . | . | . | . | . | . | G |
| F | . | . | . | . | . | . | A | . | . | . | . | . | . | . | . | T | . | . | . |
| G | . | . | . | . | . | . | A | . | . | . | . | . | . | . | . | T | . | T | . |
| H | T | A | A | G | T | G | . | C | . | T | A | G | A | G | C | T | A | T | . |

| Hap | 418 | 473 | 503 | 524 | 544 | 557 | 575 | 578 | 587 | 641 | 722 | 770 | 787 | 794 | 824 | 827 | 833 | 884 | 914 |
| --- | --- | --- | --- | --- | --- | --- | --- | --- | --- | --- | --- | --- | --- | --- | --- | --- | --- | --- | --- |
| A | A | G | A | G | G | C | G | G | T | G | A | C | A | T | G | T | A | T | T |
| B | . | . | . | . | . | . | . | . | . | . | . | . | . | . | . | . | . | . | . |
| C | . | . | . | . | . | . | . | . | . | . | . | . | . | . | . | . | . | . | . |
| D | . | . | . | . | . | . | A | . | . | . | . | . | . | . | . | . | . | . | . |
| E | . | . | . | . | . | . | A | . | . | . | . | . | . | . | . | . | . | . | . |
| F | . | . | . | . | . | . | . | . | . | . | . | . | G | . | . | C | . | . | . |
| G | . | . | . | . | . | . | . | . | . | . | . | . | G | . | . | . | . | . | . |
| H | G | A | G | A | A | T | . | A | C | A | G | T | G | C | A | . | G | C | C |

| Hap | 923 | 926 | 932 | 943 | 959 | 970 | 998 | 1001 | 1025 | 66 | 68 | 183 | 268 | 309 | 321 | 411 | 447 |
| --- | --- | --- | --- | --- | --- | --- | --- | --- | --- | --- | --- | --- | --- | --- | --- | --- | --- |
| A | T | G | A | C | G | C | T | A | T | T | G | A | T | A | C | C | C |
| B | . | . | . | T | . | . | . | . | . | . | . | . | . | . | . | . | . |
| C | . | . | . | . | . | . | . | . | . | . | . | . | . | . | . | . | . |
| D | C | . | . | . | . | . | . | . | . | . | . | G | . | . | . | . | . |
| E | . | . | . | . | . | G | . | . | C | . | . | . | . | . | . | . | . |
| F | . | . | . | . | A | . | . | . | . | . | A | G | . | . | . | T | . |
| G | . | . | . | . | A | . | . | . | . | . | A | G | . | . | . | T | . |
| H | . | A | G | . | . | . | C | G | . | C | A | G | C | C | T | . | T |

Haplotypic diversity across all of our samples was 0.55, being highest in the Lower Mississippi River population (0.60±0.05), followed by the Upper Mississippi River, and lowest in the Missouri River (0.46±0.09); these values did not significantly vary (Table 2). The two most common haplotypes, “A” and “B”, occurred throughout all of the populations (Fig. 3) and were distinguished by a single nucleotide (Table 5, Fig. 4). Most of the remaining silver carp haplotypes differed by one to six nucleotide steps from haplotype “A”, with the exception of “H” (whose divergence was much greater; Fig. 4). Haplotypes exclusively found outside of North America, cyt*b*-“R” and cyt*b*-“Q”, were closest in sequence to our rare “F” and “G” haplotypes, differing by five to nine steps from haplotype “A” (Fig. 4).

The haplotype networks revealed a distinctly divergent haplotype “H”, which occurred in all populations except for the Illinois River, and was more than 44 sequence steps from the closest silver carp haplotype and 74 steps from the bighead carp (Figs. 3–4, Table 5). The Bayesian evolutionary trees (Fig. 5) showed strong support for the respective silver carp and bighead carp clades, which each had posterior probabilities of 1.00. The “H” haplotype was placed as a basal branch outside of the silver carp clade, with 1.00 posterior probability support (Fig. 5A).

Analyses of the nuclear S7 sequences revealed 21 “fixed” nucleotide differences between the silver and bighead carp species (Fig. 5B). Individuals having the “H” mtDNA haplotype did not vary from other silver carp in their S7 intron sequences, and lacked a common genotype or pattern. Many of them shared identical sequences for the nuclear S7 intron with other “H” individuals and other silver carp haplotypes, while others differed in their S7 sequences alone (Table S2). For example, 12 individuals possessed the same S7-“1” sequence, including four having mtDNA haplotype “H”, two with haplotype “A”, and one each with haplotypes “B–G”. Individuals that possessed sequences S7-“2” and −“3” each contained one individual of haplotype “A”, whereas three individuals with S7-“4” had haplotype “B”. A silver carp having the S7-“5” sequence contained haplotype “G”, and one with the S7-“6” sequence had haplotype “H” (Table S2).

There was no nuclear haplotype genetic differentiation of silver carp individuals that possessed the highly divergent mtDNA haplotype “H” from the 14 other silver carp individuals examined, which included all other mtDNA haplotypes, according to the *θ*_ST_ and exact test analyses. Thus, the mtDNA “H” variant showed no similar divergence in nuclear DNA, in either their microsatellite allelic frequencies or S7 sequences.

### High-throughput sequencing results of eDNA from retailers

After bioinformatic filtering (including a cutoff of 0.02% of total sequences passing quality control per sample for both assays, calculated based on the positive controls), our assays discerned genetic evidence of silver carp haplotype “B” in the water from two bait shops, which were located in the Sandusky Bay and Maumee Bay regions of western Lake Erie. Silver carp was not identified from any of the morphological fish samples purchased from the 45 bait shops, in the water samples from any other shops, or in our controls. An estimated 0.2% of the sequences from the shop in the Sandusky Bay region belonged to silver carp, based on assay one results. Amplification of the sample for assay two was lower than its cutoff, but its detection was deemed valid according to our study’s detection criteria, due to the large proportion of positive reads from assay one. In the shop from the Maumee Bay region, 0.02% of the reads from assay one were identified as silver carp, and for assay two, 13.85% of the reads matched silver carp haplotype B and 0.052% were identified as a “new” (previously unknown) haplotype (Table S3). NTCs did not amplify and no silver carp detections occurred in positive controls or in the 43 other bait shop samples. For this reason and because silver carp DNA was identified using two separate assays from libraries sequenced on two different MiSeq runs, it is highly unlikely this positive detection was the result of contamination. The bait shops in this region sometimes sold bait fishes that were obtained from sources in the south in the Mississippi River system, where silver carp are resident.

We also discerned silver carp in water samples from two pond supply stores sampled in 2017, which were both located in the Lake St. Clair watershed. Both assays were positive for both stores, and both stores contained silver carp of haplotype “B”. In addition, one of the stores also contained eDNA of bighead carp, which also was confirmed with both assays.

## Discussion

### Genetic diversity patterns and the “leading edge” hypothesis

Analyses of population genetic variation across the invasive range of silver carp using microsatellites and DNA sequences elucidated relatively high levels of genetic diversity (measured by heterozygosity and allelic richness), as well as consistency in their levels across the population samples. There were no significant differences in the invasion front populations versus longer-established populations. This refutes the leading edge hypothesis of lower levels of genetic diversity for the populations of silver carp at the two invasion fronts. Moreover, significance genetic divergence was discerned between the two invasive front populations, with the microsatellite results.

Our findings indicate that silver carp populations across the North American invasion possess relatively high genetic diversity levels (*A*_R_=6.1±0.07, *H*_O_=0.60±0.14), in comparison to other invasive fish populations (see [6]). For example, allelic richness was much lower for the Eurasian ruffe invasion of the Great Lakes (*A*_R_=3.09±0.91) and heterozygosity was similar (*H*_O_=0.59±0.03), using a similar number of microsatellite loci [6]. In comparison, allelic richness for the Eurasian round goby’s invasion of the Great Lakes was much higher (*A*_R_=9.01±0.92) and heterozygosity levels were similar (*H*_O_=0.60±0.05), again based on a similar number of microsatellite loci [5]. Population genetic data for silver carp from its native range are scarce, making it difficult to directly compare levels of genetic diversity and structure to its native populations. A study by Farrington *et al.* 2017 [95] found allelic richness in the North American population at 4.7±0.1 versus 6.1±0.2 in the native range, using their microsatellite alleles, whose values for the former were lower than those discerned here. Their mean heterozygosity values appeared slightly greater (0.65) than ours. These variations likely reflect the loci used and the population samples analyzed.

Li *et al.* 2011 [96] reported 18 unique mtDNA haplotypes (based on concatenated COI and control region (D-loop) sequences) in 94 silver carp individuals sampled from the Illinois and Mississippi rivers, which were not detected in samples from their native range (*N*=121; Yangtze, Pearl, and Amur Rivers). Haplotypic diversity was reported to be lower in the Mississippi River basin (0.74) than reported in the native range (0.95) in their study. Unfortunately, their sequences are not in GenBank or in any accessible data bases, and the authors have not responded to numerous requests to share their data. Our haplotypic diversity was 0.61 in the Lower Mississippi, ranging down to 0.48 in the Missouri River, and averaged 0.54 across all of our samples. Our results and those from other studies suggest a small founder effect for the introduced silver carp in North America.

Some invasive fish populations have been successful in new areas despite reduced genetic diversity levels, in comparison to native ones. For example, the Indo-Pacific lionfishes *Pterois volitans* and *P. miles*, which are widely invasive and prolific in the Western Atlantic Ocean and the Caribbean Sea, respectively had nine haplotypes and just a single haplotype in their introduced ranges, versus 36 and 38 in their native ranges [97]. Independent, allopatric introductions of Eurasian ruffe into the upper Great Lakes and Bassenthwaite Lake, England showed significant founder effects in microsatellite allelic richness and heterozygosity levels, with little differentiation across the geographic extents of the invasions, low effective population sizes, and declines in genetic diversity over their 30-year time courses [6]. Sacramento pikeminnow *Ptychocheilus grandis* populations have been highly successful invaders into and from coastal Californian rivers despite a 49.6% reduction in allelic richness due to founding by only four individuals [98]. Chen *et al.* 2012 [99] compared introductions of grass carp on three continents to their native range, discerning reduced genetic variation in all invasive populations. Evidence for founder effect and bottlenecks was indicated for two of the locations, but not in North America. The authors hypothesized that rapid expansion into the large Mississippi River basin and likely multiple introduction events have counteracted the effects of bottlenecks on those populations. Our results implicate that similar patterns have influenced genetic diversity in silver carp populations.

In contrast, Roman and Darling 2007 [12] described that just 37% of introduced populations actually experienced a significant loss of genetic diversity during their establishment. For example, most introduced populations of wakame algae *Undaria pinnatifida* contained high genetic diversity worldwide, likely stemming from admixture of multiple native strains [100]. Likewise, the ballast water introductions of dreissenid mussels and the round goby in the Great Lakes were large and very genetically diverse, involving several likely source populations and introduction events, and showed no founder effects [5, 7, 15, 16, 18, 19]. Such ballast water introductions have involved up to hundreds or thousands of individual propagules being introduced at a time [12, 101]. These contrast with the apparently more limited diversity of the silver carp invasion.

The leading edge hypothesis predicts that species with rapidly expanding ranges may possess lower genetic diversity in the more peripheral populations [20, 102]. This hypothesis has been supported for several native species that expanded from glacial refugia [20], such as the yellow perch *Perca flavescens* in the Great Lakes [103]. The leading edge hypothesis was supported by a single-nucleotide polymorphism (SNP) analysis of three invasive fronts for the bank vole *Myodes glareolus* introductions in Ireland, which showed reduced genetic diversity, but no accumulation of deleterious alleles, and possible selection for traits involved in immunity and behavior [104]. Low diversity at the leading edge was implicated in a study of variation in North American expansion areas for the vampire bat *Desmodus rotundus* [21]. When invasive species spread via jump dispersal, new population areas often possess reduced genetic diversity, as was seen with invasive dreissenid mussels spreading from the Great Lakes to smaller lakes to the west [15, 16] and with the European green crab’s *Carcinus maenas* spread from the eastern to western coasts of North America [105].

However, the leading edge hypothesis is not supported across all invasions, such as the round goby, which showed no variations in genetic diversity levels across its introduced range in the Great Lakes and whose populations have remained genetically divergent yet stable over the 25 year course of the invasion [5]. The present study likewise discerns that silver carp populations do not significantly vary in genetic diversity, assessed with allelic richness and observed heterozygosity, across its invasive range – including at two leading edge invasion fronts. This may have resulted from their steady expansion without occurrence of jump dispersal, and the continued maintenance of large population sizes and gene flow across the introduced range. The considerable geographic distances separating three of the four pairs of identified full siblings of silver carp in our study also support the hypothesis of high dispersal, including among members of cohorts.

### Genetic divergence patterns and population relationships

Significant population genetic differentiation sometimes occurs across the range of invasive species, attributed to high diversity and multiple founding sources, including the round goby [19], whose population patterns then have remained stable over 25 years [5]. The Eurasian ruffe exhibited population divergence across its invasive range in the upper Great Lakes, revealing barriers to gene flow; these population differences have been maintained across the 30 years of the invasion [6]. In contrast, the genetic composition of the introduced zebra mussel [15, 16] has changed over the invasional time period of three decades in the Hudson River, experiencing several population turn-overs and recolonizations, yet has remained consistent in Lake Erie during this period [106]. In comparison, some invasions possess little population genetic structure across their ranges. For example, the Virginia warbler *Oreothlypis virginiae*, which recently colonized the Black Hills of South Dakota, showed lack of differentiation across its range, implicating high gene flow and dispersal [107].

In the present study, the silver carp displays modest levels of inter-population divergences across its invasive North American range, with most of the exact tests of population differentiation being significant. Population differentiation is more pronounced for the southern Mississippi River area of original introduction versus the other areas of their population expansions. Overall, the silver carp invasion appears to have progressed steadily, without bottlenecks or marked changes in population composition. Most alleles are distributed throughout the populations, with some slight, yet significant differences at the invasion fronts of the Illinois and Wabash Rivers. The latter showed high self-assignment.

### Genetic history of the invasion, revealed by mtDNA

Interpreting the genetic composition, diversity, and phylogeographic patterns of introduced populations and their sources is necessary for understanding how organisms colonize, expand their ranges, [108] and potentially adapt to their new environments [2]. The origins of North American silver carp are believed to trace to China, based based on aquaculture records summarized by Kolar *et al.* 2007 [109] and prior genetic work by Li *et al.* 2011 [96]. The latter study’s haplotype network suggested that the two highest frequency North American haplotypes found in our study likely traced to the Amur and/or Yangtze Rivers. We compared our sequences from the invasive range to those from the native range in the Yangtze River (cyt *b* haplotype “Q”) and from an introduction to the Black River in Russia (“R”), whose sequences were closest to our rare “F” and “G” haplotypes. Haplotypes “A” and “B” occurred at the highest frequencies in our study, and in similar proportions at most locations, with “A” being more common in the Wabash River invasion front and the Missouri River population, and “B” being more prevent at the Illinois River invasion front. Our “A” and “B” haploytpes likely correspond to the most common haplotypes noted by Li *et al.* 2011 [96], but since their sequences were not deposited in GenBank or in other publicly-available data bases, are not provided in their paper, and the authors have not responded to requests from us to provide them, this cannot be verified.

The singleton haplotypes “C”, “D”, and “E” discerned in our analysis, all occurred in the Lower Mississippi River population, and “F” was from the Upper Mississippi River. Individuals from the invasive front populations (Illinois and Wabash Rivers) were dominated by the two common haplotypes, except for one individual from the Illinois River that shared the “G” haplotype with a Lower Mississippi River fish. These results suggest that the original introduction(s) of silver carp into the Lower Mississippi River region founded the invasion, and that there likely were not additional introductions from other native sources into more peripheral populations.

### Introgression with largescale silver carp

A very divergent mtDNA haplotype – “H”– occurred in about 3% of the individuals that we sequenced, which may represent a historic introgression of *H. molitrix* with the related largescale silver carp species, *H. harmandi* Sauvage, 1884. Our mtDNA sequences for haplotype “H” matched a GenBank haplotype (#EU315941) collected in the Yangtze River of China. The two mtDNA sequences differed at 38 nucleotide positions (Table 5). The largescale silver carp *H. harmandi* is morphologically distinguishable in possessing larger scales and a deeper body than does *H. molitrix* [110]. The largescale silver carp is native to Hainan Island and Vietnam, where silver carp also has been extensively introduced [110]. The two species have widely hybridized in China and Vietnam [110]. The nuclear DNA S7 sequences and microsatellite allelic frequencies possessed by the mtDNA “H”-haplotype individuals that we analyzed indicate that these individuals lack nuclear DNA differentiation. The mtDNA “H” haplotype appears to have resulted from mtDNA introgression, likely of female *H. harmandi* interbreeding with male *H. molitrix* in Asia. Our analyses found no nuclear DNA differentiation for those individuals, in either microsatellite alleles or S7 sequences. Our findings discern that the *H. harmandi* mtDNA haplotype “H” is widespread throughout the invasive North American range, occurring in our samples from the Lower Mississippi, Upper Mississippi, Missouri, and Illinois Rivers, as well as in China.

Compared to Farrington *et al.* 2017 [95], who sequenced the entire mitochondrion of 30 silver carp individuals from North America and two from their native range, we sequenced 225 individuals for ~10% of some of the most variable portiion of the tmitochondrial genome, finding eight haplotypes compared to their six, including a much higher frequency of the highly divergent “H” haplotype (for which they uncovered just a single individual) (Table 5). We included Farringon et al.’s 2017 [95] sequences in our analyes, except for four of their haplotypes, which either were outside of the cyt *b* and COI gene that we sequenced, or had incomplete sequences. With our greater sample size, we discerned that the “H” haplotype occurs throughout most of the North American range. Farrington et al. 2017 [95] found a single individual with the “H” haplotype (which was >415 nt steps from the most similar haplotype) in the Upper Mississippi River alone.

Our S7 nuclear gene tree showed congruent relationships with the mtDNA tree, except that S7 sequences of the highly divergent “H” haploytpe did not differ from other silver carp. Introgressed native silver carp were likely present as a small percentage of founders in the North American introduction.

Wild silver and bighead carp likely do not widely interbreed in their native range, having developed pre-zygotic reproductive barriers in sympatry [109]. Genetic studies have demonstrated that the two species often hybridize in aquaculture [57] and hybrids occur across their invasive North American range [33, 34]. Lamer *et al.* 2015 [34] used 57 nuclear single nucleotide polymorphisms (SNPs) and one mtDNA polymorphism to determine that 44.7% of *Hypophthalmichthys spp*. in the Mississippi River Basin have interbred or backcrossed, suggestive of a hybrid swarm. The flow of maternal DNA was found to be biased from silver carp to bighead carp in both the F1’s and bighead carp backcrosses. Across all of our samples, we identified that just a single individual had bighead carp mtDNA, either due to misidentification or hybridization. There are 73 bp changes separating haplotype “H” from the most similar bighead carp cyt *b* sequence. The silver carp haplotype “Q”, which is most similar to our “H”, diverges from it by 37 bp; therefore, it is highly unlikely that haplotype “H” was the result of hybridization with bighead carp. Given our objective to describe the mtDNA phylogeographic pattern of silver carp in the invasive range, it may be important to consider how the interaction between the two species may influence that pattern in the future. It is possible that the initial and ongoing hybridization acted to enhance invasive success by alleviating some of the small loss of additive genetic variation experienced with the founding events.

### eDNA detection in retail trade

We did not physically find silver carp in any of the 45 bait shops or the 22 pond supply stores surveyed, based on morphological sampling. However, we discovered eDNA in the tanks of two bait shops and two pond supply retailers, where silver carp likely were present at low densities. In addition, eDNA of bighead carp was identified from one of those pond supply stores. Our eDNA findings highlight the probability that silver and bighead carps are prevalent in the retail trade in the Great Lakes watershed, where they likely blend in with other “minnows” for sale as bait. Releases from bait and pond stores may comprise an important vector for their introduction.

The eDNA HTS assays to detect their presence, as well as others in development by our laboratory, identify invasive carps in water samples to species and also discern their population structure. Other published assays that detect invasive carps have relied on qPCR [111], cannot differentiate haplotypes, and therefore do not provide any information on genetic diversity within or among populations. The “new” haplotype found here was not recovered in our traditional Sanger sequencing of silver carp from throughout its invasive range and is not referenced in GenBank. Despite our use of a de-noising algorithm and error cutoffs calculated using positive controls, this previously undetected haplotype might have stemmed from sequencing error introduced from the Illumina MiSeq platform, possibly originating from haplotype “B”. Further refining of bioinformatic pipelines and the use of PCR replicates could improve confidence in the presence or absence of rare “new” haplotypes discovered with eDNA assays.

Our diagnostic HTS assays have potential widespread use in screening of fish sold by bait and pond store retailers, as well as in water and ichthyoplankton samples from the environment. The latter will allow early identification of eggs and larvae to species, which are presently impossible to morphologically discern. Moreover, screening plankton samples using these markers will allow relative species proportions to be evaluated, as well as yield population genetic variability statistics (see [106]) for ecological comparisons.

### Conclusions

The present investigation resolved phylogeographic and genetic diversity patterns across the invasive native range of silver carp, significantly advancing from prior knowledge using larger sample sizes and a combined population genetic approach. We discerned relatively high levels of genetic diversity across its invasive North American range, including at the fronts leading to the Great Lakes. The latter refuted the leading edge hypothesis for the silver carp, indicating that the invasion has maintained high numbers of diverse individuals in colonizing areas. This also was supported by the occurrence of widely-dispersed full siblings. We discovered a widespread and highly differentiated ancient mtDNA haplotype, which likely originated from historic introgression with the largescale silver carp in Asia, predating the North American introduction. The introgressed individuals do not differ from silver carp in nuclear DNA microsatellites or gene sequences, suggesting that female silver carp reproduced with male largescale silver carp.

There is great interest in preventing entry of silver carp into the Great Lakes system with man-made barriers and the use of genetic tools to detect them at the leading edges or in other possible introduction vectors. Our finding of eDNA from silver carp in two bait stores in the Lake Erie watershed points to another likely vector for introduction, as the bait stores often source bait from southern states in the Mississippi River watershed, where silver carp are well-established.

Most other eDNA detection approaches are based on quantitative PCR (qPCR) analyses for detecting presence or absence of a given single target species. Our eDNA assay is designed to discern and detect multiple divergent haplotypes, including the introgressed haplotype “H”, as well as distinguish among related cyprinid species. It can be used on water samples or with ichthyoplankton tows, which will yield abundant population genetic information. At early life stages, there are few or no characters to distinguish among these cyprinid species. The ability to track shifts in species composition and population genetic variation, rather than just single species presence or absence using eDNA techniques may enhance our understanding of how invasive carps have been so successful in colonizing new areas. If silver carp spread into the Great Lakes via natural dispersal, populations likely will possess high genetic diversity, yet may subtly vary in genetic composition, allowing tracking. The fact that the two invasive fronts analyzed here differ in genetic composition, as well as from the longer-established core area of the lower Mississippi River, shows the influence of genetic drift and possibly adaptation. Elucidating these genetic patterns will significantly increase ecological understanding of the relative successes and adaptations of invasions, and aid the rapid identification of new populations and range areas.

## Acknowledgments

This work was funded by a grant award to CAS from the USEPA Great Lakes Restoration Initiative (GLRI) #GL-00E01898. Specimens either were collected by us and other members of CAS’ Genetics and Genomics Group laboratory or were provided by various collectors, including: M. Bartron, D. Chapman, A. Coulter, J. Lamar, M. Pyron, and R. Lance. We thank F. Calzonetti and S. McBride for logistical help. This is NOAA Pacific Marine Environmental Laboratory (PMEL) contribution #XXX.

## Author Contributions

Conceptualization: CAS.

Data curation: AEE, MRS, CAS.

Formal analysis: AEE, MRS, CAS.

Funding acquisition: CAS.

Investigation: CAS.

Methodology: CAS.

Project administration: CAS.

Resources: CAS.

Supervision: CAS.

Validation: CAS.

Visualization: CAS, AEE, MRS.

Writing ± original draft: CAS, AEE, MRS.

Writing ± review & editing: CAS, AEE, MRS.

---

**Supplementary Table 1.**
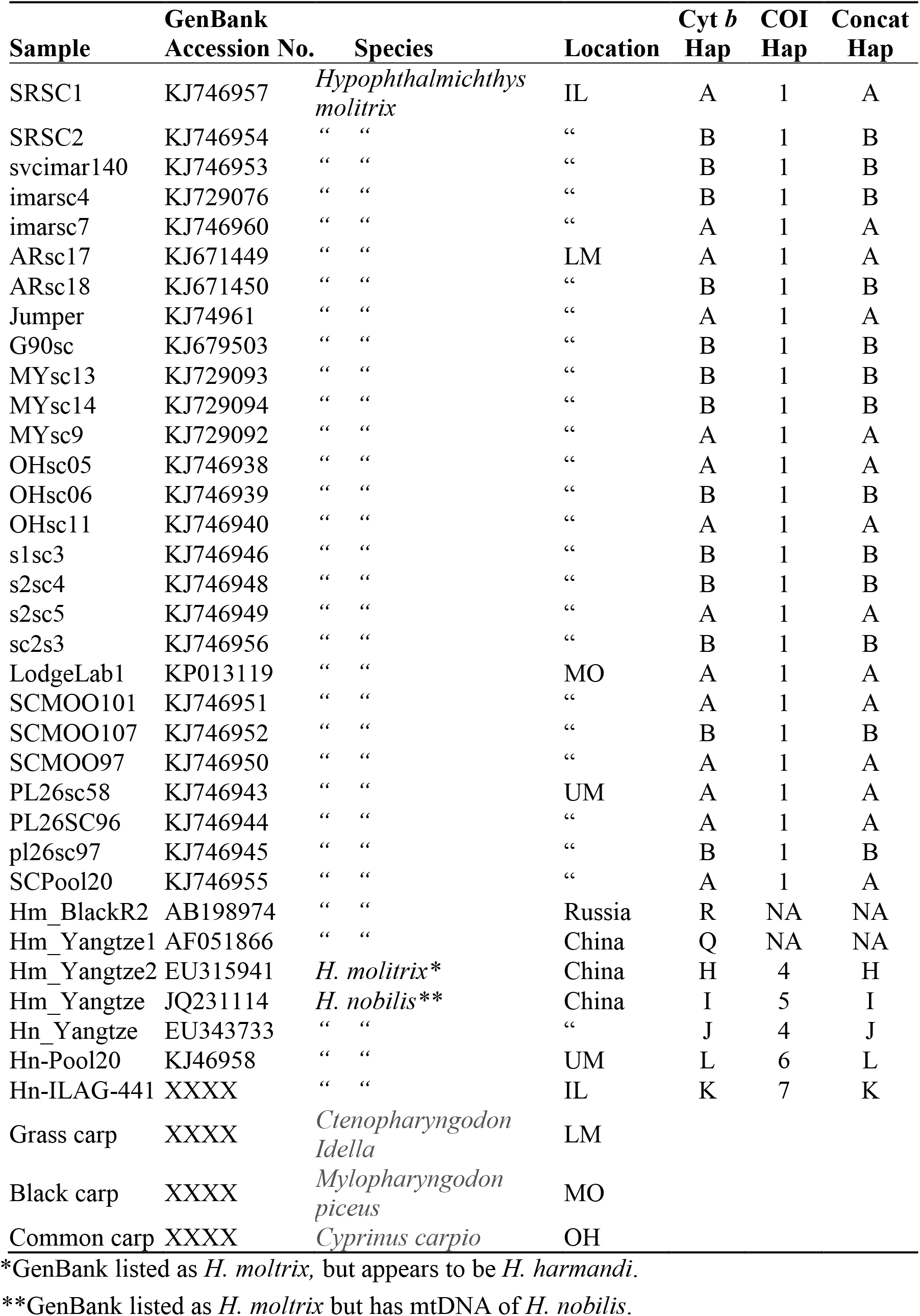
Samples, GenBank accessions, and their haplotypes (Hap), which are included in our mtDNA sequence analyses.

**Supplementary Table 2.** S7 nuclear ribosomal protein gene, intron 1, partial sequence haplotypes of silver carp.

| **Hap** | 150 | 202 | 253 | 273 | 403 | 445 | 462 | 468 | 558 | 577 | 646 | 647 | 648 | 649 | 650 | 658 | 661 | 669 | 732 | 733 | 735 |
| --- | --- | --- | --- | --- | --- | --- | --- | --- | --- | --- | --- | --- | --- | --- | --- | --- | --- | --- | --- | --- | --- |
| **S7-1** | T | C | A | C | G | T | C | G | T | A | - | - | - | C | C | T | C | T | - | - | C |
| **S7-2** | . | T | . | . | . | C | . | A | . | . | - | - | - | - | - | . | . | . | A | C | G |
| **S7-3** | . | . | . | S | R | . | M | . | . | . | - | - | C | C | C | C | . | . | W | C | S |
| **S7-4** | . | . | . | . | . | . | . | . | . | . | - | - | C | C | C | . | . | . | - | - | . |
| **S7-5** | A | T | G | . | A | . | A | . | C | G | - | - | - | C | C | . | T | C | - | - | . |
| **S7-6** | . | . | G | G | A | . | A | . | C | G | C | C | C | C | C | . | T | C | - | - | . |

Nucleotide positions (above sequence) based on alignment to GenBank accession AY325777 reference sequence. Dots indicate a base identical to haplotype “A-1”. Dashes denote a deletion in the sequence. Only variable bases are shown.

**Supplementary Table 3.** Silver carp haplotypes discerned with our eDNA high throughput sequencing assay from a water sample from a bait shop in the western Lake Erie watershed. Table includes number of sequences (*N* seqs), and percent of total sequences passing quality control in the sample (% total seqs). Nucleotide positions (above sequence) based on complete cytochrome *b* gene. Only the sole variable nucleotide differing between haplotype B and the “new” haplotype is shown.

| **Hap** | ***N* Seqs** | **% total seqs** | **87** |
| --- | --- | --- | --- |
| **B** | 6736 | 13.85 | A |
| **“new”** | 254 | 0.52 | C |

## References

1. Mooney H, Cleland E. The evolutionary impact of invasive species. Proc Natl Acad Sci USA. 2001; 98:5446–5451. https://doi.org/10.1073/pnas.091093398

2. Wares JP, Hughes AR, Grosberg RK. Mechanisms that drive evolutionary change: insights from species introductions and invasions. In: Sax DF, Stachowicz JJ, Gaines SD, editors. Species Invasions: Insights into Ecology, Evolution, and Biogeography. Sunderland, Massachusetts: Sinauer Associates; 2005. pp. 229–257.

3. Colautti, RI, Lau JA. Contemporary evolution during invasion: evidence for differentiation, natural selection, and local adaptation. Mol Ecol. 2015; 24:1999–2017. https://doi.org/10.1111/mec.13162

4. Huey RB, Gilchrist GW, Hendry AP. Using invasive species to study evolution: case studies with Drosophila and salmon. In: Sax DF, Stachowicz JJ, Gaines SD, editors. Species Invasions: Insights into Ecology, Evolution, and Biogeography. Sunderland, Massachusetts: Sinauer Associates; 2005. pp. 139–164.

5. Snyder MR, Stepien CA. Genetic patterns across an invasion's history: a test of change versus stasis for the Eurasian round goby in North America. Mol Ecol. 2017; 26:1075–1090. https://doi.org/10.1111/mec.13997 PMID: 28029720

6. Stepien CA, Eddins D, Snyder M, Marshall NT. Genetic change versus stasis over the time course of invasions: trajectories of two concurrent, allopatric introductions of the Eurasian ruffe. Aquat Invasions. Accepted. 2018

7. Stepien CA, Brown JE, Neilson ME, Tumeo MA. Genetic diversity of invasive species in the Great Lakes versus their Eurasian source populations: insights for risk analysis. Risk Anal. 2005; 25:1043–1060. https://doi.org/10.1111/j.1539-6924.2005.00655.x PMID: 16268948

8. Forsman A. Effects of genotypic and phenotypic variation on establishment are important for conservation, invasion, and infection biology. Proc Natl Acad Sci USA. 2014; 111:302–307. https://doi.org/10.1073/pnas.1317745111 PMID: 24367109

9. Baker HG, Stebbins GL. The Genetics of Colonizing Species. New York, New York: Academic Press; 1965.

10. Williamson M. Invasions. Ecography. 1999; 22:5–12. https://doi.org/10.1111/j.1600-0587.1999.tb00449.x

11. Bock DG, Caseys C, Cousens RD, Hanh MA, Heredia SM, Hubner S, Turner KG, Whitney KD, Risenberg LH. What we still don’t know about invasion genetics. Mol Ecol. 2015; 24:2277–2297. http://onlinelibrary.wiley.com/doi/10.1111/mec.13032/full

12. Roman J, Darling JA. Paradox lost: genetic diversity and the success of aquatic invasions. Trends Ecol Evol. 2007; 22:454–464. https://doi.org/10.1016/j.tree.2007.07.0021002/aqc.2414

13. Lockwood JL, Hoopes MF, Marchetti MP. Propagules. In: Invasion Ecology, 2nd Edition. West Sussex, UK: Wiley-Blackwell; 2013. pp. 75–98.

14. Rius M, Turon X, Bernardi G, Volckaert FAM, Viard F. Marine invasion genetics: from spatio-temporal patterns to evolutionary outcomes. Biol Invasions. 2015; 17:869–885. https://doi.org/10.1007/s10530-014-0792-0

15. Brown JE, Stepien CA. Population genetic history of the dreissenid mussel invasions: expansion patterns across North America. Biol Invasions. 2010; 12:3687–3710. https://doi.org/10.1007/s10530-010-9763-2

16. Stepien CA, Grigorovich IA, Gray MA, Sullivan TJ, Yerga-Woolwine S, Kalayci G. Evolutionary, biogeographic, and population genetic relationships of dreissenid mussels, with revision of component taxa. In: Nalepa TF, Schloesser DW, editors. Quagga and Zebra Mussels: Biology, Impacts, and Control. New York: CRC Press; 2013. pp. 403–444.

17. Stepien CA, Tumeo MA. Invasion genetics of Ponto-Caspian gobies in the Great Lakes: A “cryptic” species, absence of founder effects, and comparative risk analysis. Biol Invasions. 2006; 8:61–78. https://doi.org/10.1007/s10530-005-0237-x

18. Brown JE, Stepien CA. Ancient divisions, recent expansions: phylogeography and population genetics of the round goby Apollonia melanostoma across Eurasia. Mol Ecol. 2008; 17:2598–2615. https://doi.org/10.1111/j.1365-294X.2008.03777.x

19. Brown JE, Stepien CA. Invasion genetics of the Eurasian round goby in North America: tracing sources and spread patterns. Mol Ecol. 2009; 18:64–79. https://doi.org/10.1111/j.1365-294X.2008.04014.x

20. Hewitt GM. Some genetic consequences of ice ages, and their role in the divergence and speciation. Biol J Linnean Soc. 1996; 58:247–276. https://doi.org/10.1111/j.1095-8312.1996.tb01434.x

21. Piaggio AJ, Russell AL, Osorio IA, Jiménez Ramírez A, Fischer JW, Neuwald JL, et al. Genetic demography at the leading edge of the distribution of a rabies virus vector. Ecol Evol. 2017; 7:5343–5351. https://doi.org/10.1002/ece3.3087

22. Ochocki BM, Miller TEX. Rapid evolution of dispersal ability makes biological invasions faster and more variable. Nature Communications. 2017, 8, 14315 https://doi.org/10.1038/ncomms14315

23. Shine, R., Brown, G. P. & Phillips, B. L. An evolutionary process that assembles phenotypes through space rather than through time. Proc Natl Acad Sci USA. 2011; 108:5708–5711. https://doi.org/10.1073/pnas.1018989108

24. Perkins TA, Phillips BL, Baskett ML, Hastings A. Evolution of dispersal and life history interact to drive accelerating spread of an invasive species. Ecol Lett. 2013; 16:1079–1087. https://doi.org/10.1111/ele.12136

25. May B, Marsden JE. Genetic identification and implications of another invasive species of dreissenid mussel in the Great Lakes. Can J Fish Aquat Sci. 1992; 49:1501–1506. https://doi.org/10.1139/f92-166

26. Karatayev AY, Mastitsky SE, Padilla DK, Burlakova LE, Hajduk MM. Differences in growth and survivorship of zebra and quagga mussels: size matters. Hydrobiologia. 2011; 668:183–194. https://doi.org/10.1007/s10750-010-0533-z

27. Quinn A, Gallardo B, Aldridge DC. Quantifying the ecological niche overlap between two interacting invasive species: the zebra mussel (Dreissena polymorpha) and the quagga mussel (Dreissena rostriformis bugensis). Aquat Conserv: Mar and Freshw Ecosyst. 2014; 24:324–337. https://doi.org/10.1002/aqc.2414

28. Nalepa TF, Fanslow DL, Lang GA. Transformation of the offshore benthic community in Lake Michigan: recent shift from the native amphipod Diporeia spp. to the invasive mussel Dreissena rostriformis bugensis. Freshwater Biol. 2009 54, 466–479. https://doi.org/10.1111/j.1365-2427.2008.02123.x

29. Nalepa, TF, Fanslow DL, Pothovern SA. Recent changes in density, biomass, recruitment, size structure, and nutritional state of Dreissena populations in southern Lake Michigan. J Great Lakes Res. 2010; 36 (Supple. 3):5–19. https://doi.org/10.1016/jjglr.2010.03.013

30. Jude DJ, Reider RH, Smith GR. Establishment of Gobiidae in the Great Lakes basin. Can J Fish Aquat Sci. 1992; 49:416–421. https://doi.org/10.1139/f92-047

31. Kocovsky, P.M., J.A. Tallman, D.J. Jude, D.M. Murphy, J.E. Brown & C. A. Stepien. 2011. Expansion of tubenose gobies Proterorhinus semilunaris into western Lake Erie and potential effects on native species. Biol Invasions. 13(12):2775–2784. https://doi.org/10.1007/s10530-011-9962-5

32. Freeze M, Henderson S. Distribution and status of the bighead carp and silver carp in Arkansas. North Am J Fish Mana. 1982; 2:197–200. https://doi.org/10.1577/1548-8659(1982)2<197:DAS0TB>2.0.CO;2

33. Lamer JT, Dolan CR, Peterson JL, Chick JH, Epifanio JM. Introgressive hybridization between Bighead Carp and Silver Carp in the Mississippi and Illinois Rivers. North Am J Fish Mana. 2010; 30:1452–1461. https://doi.org/10.1577/M10-053.1

34. Lamer JT, Ruebush BC, Arbieva ZH, McClelland MA, Epifanio JM, Sass GG. Diagnostic SNPs reveal widespread introgressive hybridization between introduced Bighead and Silver Carp in the Mississippi River Basin. Mol Ecol. 2015; 24:3931–3943. https://doi.org/10.1111/mec.13285 PMID:26096550

35. Robison HW, Buchanan TM. Fishes of Arkansas. Fayetteville, AR: University of Arkansas Press; 1988.

36. Koel TM, Irons KS, Ratcliff E. Asian carp invasion of the Upper Mississippi River System. USGS Upper Midwest Environmental Sciences Center Project Status Report. 2000. Available from: https://umesc-usgs-gov.proxy.lib.umich.edu/documents/project_status_reports/2000/psr00_05.pdf

37. Cooke SL, Hill WR. Can filter-feeding Asian carp invade the Laurentian Great Lakes? A bioenergetic modeling exercise. Freshw Biol. 2010; 55:2138–2152. https://doi.org/10.1111/j.1365-2427.2010.02474.x

38. Cudmore B, Mandrak NE. Assessing the biological risk of Asian carps to Canada, pp 15–30 in: Chapman DC, Hoff MH (eds). Invasive Asian carps in North America. American Fisheries Society Symposium 74, American Fisheries Society, Bethesda, MD. 2011.

39. Ricciardi A. Facilitative interactions among aquatic invaders: is an “invasional meltdown” occurring in the Great Lakes? Can J Fish Aquat Sci. 2001; 58:2513–2525. https://doi.org/10.1139/cjfas-58-12-2513

40. Embke HS, Kocovsky PM, Richter CA, Pritt JJ, Mayer CM, Qian SS. First direct confirmation of grass carp spawning in a Great Lakes tributary. J Great Lakes Res. 2016; 42:899–903. https://doi.org/10.1016/jjglr.2016.05.002

41. Naylor R, Williams S, Strong DR. Aquaculture—a gateway for exotic species. Science 2001; 294:1655–1656. https://doi.org/10.1126/science.1064875

42. Canonico GC, Arthington A, McCrary JK, Thieme ML. The effects of introduced tilapias on native biodiversity. Aquatic Conserv: Mar Freshw Ecosyst. 2005; 15:463–483. https://doi.org/10.1002/aqc.699.

42a. J Shellfish Res. 2007; 26:281–294. https://doi-org.proxy.lib.umich.edu/10.2983/0730-8000(2007)26[281:BAAESA]2.0.œ2

43. McKindsey CW, Landry T, O’Beirn FX, Davies IM. Bivalve aquaculture and exotic species: a review of ecological and considerations and management issues. J Shellfish Res. 2007; 26:281–294. https://doi.org/10.2983/0730-8000(2007)26[281:BAAESA]2.0.CO;2

44. Irons KS, Sass GG, McClalland, Stafford JD. Reduced condition factor of two native fish species coincident with the invasion of non-native Asian carps in the Illinois Rover, U.S.A. Is this evidence for competition and reduced fitness? J Fish Biol. 2007; 71:258–273. https://doi.org/10.1111/j.1095-8649.2007.01670.x

45. Sampson SJ, Chick JH, Pegg MA. Diet overlap among two Asian carp and three native fishes in backwater lakes on the Illinois and Mississippi Rivers. Biol Invasions. 2009; 11:483–486. https://doi.org/10.1007/s10530-008-9265-7

46. Sass GG, Hinz C, Erickson AC, McCLelland NN, McClelland MA, Epifanio JM. Invasive bighead and silver carp effects on zooplankton communities in the Illinois River, Illinois, USA. J Great Lakes Res. 2014; 40:911–921. https://doi.org/10.1016/j.jglr.2014.08.010

47. DeBoer JA, Anderson AM, Casper AF. Multi-trophic response to invasive silver carp (Hypophthalmichthys molitrix) in a large floodplain river. Freshwater Biol. 2018; 63:597–611. https://doi.org/10.1111/fwb.13097

48. Irons KS, Delain SA, Gittinger E, Ickes BS, Kolar CS, Ostendorf D, et al. Nonnative Fishes in the Upper Mississippi River System. Reston, VA: USGS; 2009. Available from: https://pubs.usgs.gov/sir/2009/5176/pdf/sir2009-5176.pdf

49. Nico L, Fuller P, Li J. Hypophthalmichthys molitrix (Valenciennes in Cuvier and Valenciennes, 1844). U.S. Geological Survey, Nonindigenous Aquatic Species Database. 2015. Available from: https://nas.er.usgs.gov/queries/FactSheet.aspx?speciesID=549

50. Coulter AA, Keller D, Amberg JJ, Bailey EJ, Goforth RR. Phenotypic plasticity in the spawning traits of bigheaded carp (Hypophthalmichthys spp.) in novel ecosystems. Freshwater B. 2013; 58:1029–1037. https://doi.org/10.1111/fwb.12106

51. Cuddington K, Currie WJS, Koops MA. Could an Asian carp population establish in the Great Lakes from a small introduction? Biol Invasions. 2014; 16:903–917. https://doi.org/10.1007/s10530-013-0547-3

52. USGS 2018 (silver carp range map) https://nas.er.usgs.gov/queries/factsheet.aspx?speciesID=549

53. Parker AD, Glover DC, Finney ST, Rogers PB, Stewart JG, Simmonds Jr. RL. Direct observations of fish incapacitation rates at a large electrical fish barrier in the Chicago Sanitary and Ship Canal. J Great Lakes Res. 2014; 41:396–404. https://doi.org/10.1016/j.jglr.2015.03.004

54. Indiana Department of Natural Resources (IDNR). Eagle Marsh berm blocks Asian carp path to Great Lakes. 12 May 2015. Available from: http://www.in.gov/activecalendar_dnr/EventList.aspx?fromdate=1/1/2017&todate=1/31/2017&display=Month&type=public&eventidn=9364&view=EventDetails&information_id=19848&print=print

55. Stern CV, Upton HR, Broughter C. 2014. Asian Carp and the Great Lakes Region. Congressional Research Service. 7–5700, R41082. Available from: https://fas.org/sgp/crs/misc/R41082.pdf

56. Great Lakes Fishery Commission (GLFC). Asian Carp. 2018. Available from: http://www.glfc.org/asian-carp.php

57. Mia MY, Taggart J, Gilmour AE, Gheyas AA, Das TK, Kohinoor AHM, et al. Detection of hybridization between Chinese carp species (Hypophthalmichthys molitrix and Aristichthys nobilis) in hatchery broodstock in Bangladesh using DNA microsatellite loci. Aquaculture. 2005; 247:267–273. https://doi.org/10.1016/j.aquaculture.2005.02.018

58. Cheng L, Liu L, Yu X, Tong J. Sixteen polymorphic microsatellites in bighead carp (Aristichthys nobilis) and cross-amplification in silver carp (Hypophthalmichthys moltrix). Mol Ecol Res. 2007; 8:656–658. https://doi.org/10.1111/j.1471-8286.2007.02037.x

59. King TL, Eackles MS, Chapman, DC. Tools for assessing kinship, population structure, phylogeography, and interspecific hybridization in Asian carps invasive to the Mississippi River, USA: isolation and characterization of novel tetranucleotide microsatellite DNA loci in silver carp Hypophthalmichthys molitrix. Cons Gen Res. 2011; 3:397–401. https://doi.org/10.1007/s12686-010-9285-3

60. Song CB, Near T, Page LM. Phylogenetic relationships among Percid fishes from mitochondrial cytochrome b DNA sequence data. Mol Phylogenet Evol. 1998; 10:343–353. https://doi.org/10.1006/mpev.1998.0542

61. Akihito T, Iwata A, Kobayashi T, Ikeo K, Ono H, Umehara Y, et al. Evolutionary aspects of gobioid fishes based upon a phylogenetic analysis of mitochondrial cytochrome b genes. Gene. 2000; 259:5–15. https://doi.org/10.1016/S0378-1119(00)00488-1

62. Ivanova NV, Zemlak TS, Hanner RH, Herbert PDN. Universal primer cocktails for fish barcoding. Mol Ecol Notes. 2007; 7:544–548. https://doi.org/10.1111/j.1471-8286.2007.01748.x

63. Chow S, Hazama K. Universal PCR primers for S7 ribosomal protein gene introns in fish. Mol Ecol. 1998; 7:1255. PMID: 9734083

64. Klymus KE, Marshall NT, Stepien CA. Environmental DNA (eDNA) metabarcoding assays to detect invasive invertebrate species in the Great Lakes. PLoS One. 2017; 12(5):e0177643. https://doi.org/10.1371/journal.pone.0177643

65. Rousset F. Genepop’008: a complete re-implementation of the GENEPOP software for Windows and Linux. Molecular Ecology Resources. 2008; 8(1):103–106. https://doi.org/10.1111/j.1471-8286.2007.01931.x

66. Zar JH. Biostatistical Analysis. 5th edition. Upper Saddle River, NJ: Prentice Hall; 2010.

67. Van Oosterhout C, Hutchinson WF, Wills DP, Shipley P. Micro-Checker: software for identifying and correcting genotyping errors in microsatellite data. Mol Ecol Notes. 2004; 4:535–538, https://doi.org/10.1111/j.1471-8286.2004.00684.x

68. Goudet J. FSTAT (Version 1.2): a computer program to calculate F-statistics. J Hered. 1995; 86(6):485–486. https://doi.org/10.1093/oxfordjournals.jhered.a111627

69. Rice WR. Analyzing tables of statistical tests. Evolution. 1989; 43:223–225. http://www.jstor.org/stable/2409177

70. R Development Core Team. R: a language for statistical computing. Version 3.2.1. R Foundation for Statistical Computing, Vienna, Austria. 2015. Available: http://www.R-project.org

71. Glaubitz JC. CONVERT: a user-friendly program to reformat diploid genotypic data for commonly used population genetic software packages. Mol Ecol Res. 2004; 4:309–310. https://doi.org/10.1111/j.1471-8286.2004.00597.x

72. Szpiech ZA, Jakobsson M, Rosenberg NA. ADZE: a rarefaction approach for counting alleles private to combinations of populations. Bioinformatics. 2008; 24(21):2498–2504. https://doi.org/10.1093/bioinformatics/btn478.

73. Jones OR, Wang J. COLONY: a program for parentage and sibship inference from multilocus genotype data. Mol Ecol Res. 2010; 10:551–555. https://doi.org/10.1111/j.1755-0998.2009.02787.x

74. Beaumont MA, Nichols RA. Evaluating loci for use in the genetic analysis of population structure. Proc R Soc Lond B. 1996; 263:1619–1626. https://doi.org/10.1098/rspb.1996.0237

75. Antao T, Lopes A, Lopes RJ, Beja-Pereira A, Luikart G. LOSITAN: A workbench to detect molecular adaptation based on a *F*_ST_-outlier method. BMC Bioinformatics. 2008; 9:323. http://www.biomedcentral.com/1471-2105/9/323

76. Weir BS, Cockerham CC. Estimating *F*-statistics for the analysis of population structure. Evolution. 1984; 38:1358–1370, https://doi.org/10.1111/j.1558-5646.1984.tb05657.x

77. Cockerham CC, Weir BS. Estimation of gene flow from F-statistics. Evolution. 1993; 43:855–863. https://doi.org/10.1111/j.1558-5646.1993.tb01239.x

78. Waples RS. Separating the wheat from the chaff: patterns of genetic differentiation in high gene flow species. J Hered. 1998; 89:438–450. https://doi.org/10.1093/jhered/89.5.438

79. Meirmans PG, Hedrick PW. Assessing population structure: *F*_ST_ and related measures. Mol Ecol Res. 2011; 11:5–18. https://doi.org/10.1111/j.1755-0998.2010.02927.x

80. Raymond M, Rousset R. An exact test for population differentiation. Evolution. 1995; 49:1280–1283. https://doi.org/10.2307/2410454

81. Narum SR. Beyond Bonferroni: less conservative analyses for conservation genetics. Conserv Genet. 2006; 7:783–787. https://doi.org/10.1007/s10592-005-9056-y

82. Benzecri JP. Statistical problems and geometric methods. Cah Anal Donnbes. 1978; 3:131–46. https://link-springer-com.proxy.lib.umich.edu/content/pdf/10.1007%2Fs10592-005-9056-y.pdf

83. Belkhir K, Borsa P, Chikhi L, Raufaste N, Bonhomme F. Genetix 4.05, logiciel sous Windows TM pour la génétique des populations. Laboratoire génome, populations, interactions, CNRS UMR 5000: 1996. Available from: http://kimura.univ-montp2.fr/genetix/

84. Pritchard JK, Stephens M, Donnelly P. Inference of population structure using multilocus genotype data. Genetics. 2000; 155:945–959. PMID: 10835412

85. Evanno G, Regnaut S, Goudet J. Detecting the number of clusters of individuals using the software STRUCTURE: a simulation study. Mol Ecol. 2005; 14:2611–2620. https://doi.org/10.1111/j.1365-294X.2005.02553.x

86. Earl DA, vonHoldt BM. Structure Harvester: a website and program for visualizing structure output and implementing the Evanno method. Conserv Genet Res. 2012; 4:359–361. https://doi.org/10.1007/s12686-011-9548-7

87. Piry S, Alapetite A, Cornuet JM, Paetkau K, Baudouin L, Estoup A. GENECLASS2: a software for genetic assignment and first generation migrant detection. J Hered. 2004; 95:536–539. https://doi.org/10.1093/jhered/esh074

88. Paetkau D, Slade R, Burden M, Estoup A. Genetic assignments for the direct, real-time estimation of migration rate: a simulation-based exploration of accuracy and power. Mol Ecol. 2004; 13:55–65. https://doi.org/10.1046/j.1365-294X.2004.02008.x

89. Clement M, Posada D, Crandall KA. TCS: a computer program to estimate gene genealogies. Mol Ecol. 2000; 9:1657–1659. https://doi.org/10.1046/j.1365-294x.2000.01020.x

90. Excoffier L and Lischer HEL. ARLEQUIN suite ver 3.5: a new series of programs to perform population genetics analyses under Linux and Windows. Mol Ecol Res. 2010; 10:564–567. https://doi.org/10.1111/j.1755-0998.2010.02847.x

91. Ronquist F, Huelsenbeck JP. MRBAYES 3: Bayesian phylogenetic inference under mixed models. Bioinformatics. 2003; 19:1572–1574. https://doi.org/10.1093/bioinformatics/btg180

92. Callahan BJ, McMurdie PJ, Rosen MJ, Han AW, Johnson AJA, Holmes SP. DADA2: high-resolution sample inference from Illumina amplicons data. Nat Methods. 2016; 13:581–583. https://doi.org/10.1038/NMETH.3869

93. Hubbs CL, Lagler KF. Fishes of the Great Lakes Region. Revised edition. Ann Arbor, MI: University of Michigan Press; 2007.

94. USGS. Nonindigenous Aquatic Species Database. 2018. Available from: https://nas.er.usgs.gov/

95. Farrington HL, Edwards CE, Bartron M, Lance RF. Phylogeography and population genetic structure of introduced silver carp (Hypophthalmichthys molitrix) and bighead carp (H. nobilis) in North America. Biol Invasions. 2017; 19:2789–2811. https://doi.org/10.1007/s10530-017-1484-3

96. Li S, Wei J, Yang Q. Significant genetic differentiation between native and introduced silver carp (Hypopthalmichtys moltrix) inferred from mtDNA analysis. Environ Biol Fish. 2011; 92:503–511. https://doi.org/10.1007/s10641-011-9870-7

97. Betancur-R. R, Hines A, Acero P, Ortí G, Wilbur AE, Freshwater DW. Reconstructing the lionfish invasion: insights into Greater Caribbean biogeography. J Biogeogr. 2011; 38:1281–1293. https://doi.org/10.1111/j.1365-2699.2011.02496.x

98. Kinziger AP, Nakamoto RJ, Harvey BC. Local-scale invasion pathways and small founder numbers in introduced Sacramento pikeminnow (Ptychocheilus grandis). Conserv Genet. 2014; 15:1–9. https://doi.org/10.1007/s10592-013-0516-5

99. Chen Q, Wang C, Lu G et al. Microsatellite genetic diversity and differentiation of native and introduced grass carp in three continents. Genetica. 2012; 140:115–123. https://doi.org/10.1007/s10709-012-9663-8

100. Voisin M, Engel CR, Viard F. Differential shuffling of native genetic diversity across introduced regions in a brown alga: aquaculture vs. maritime traffic effects. Proc Natl Acad Sci USA. 2005; 102:5432–5437. https://doi.org/10.1073/pnas.0501754102

101. Simberloff D. The role of propagule pressure in biological invasions. Annu Rev Ecol Evol Syst. 2009 40:81–102. https://doi.org/10.1146/annurev.ecolsys.110308.120304

102. Slatkin M, Excoffier L. Serial founder effects during range expansion: a spatial analog of genetic drift. Genetics. 2012; 191:171–181. https://doi.org/10.1534/genetics.112.139022

103. Sepulveda-Villet OJ & Stepien CA. Waterscape genetics of the yellow perch (Perca flavescens): patterns across large connected ecosystems and isolated relict populations. Mol Ecol. 2012; 21:5795–5826. https://doi.org/10.0000/mec.12044

104. White TA, Perkins SE, Heckel G, Searle JB. Adaptive evolution during an ongoing range expansion: the invasive bank vole (Myodes glareolus) in Ireland. Mol Ecol. 2013; 22:2971–2985. https://doi.org/10.1111/mec.12343

105. Tepolt CK, Palumbi SR. Transcriptome sequencing reveals both neutral and adaptive genome dynamics in a marine invader. Mol Ecol. 2015; 24:4145–4158. https://doi.org/10.1111/mec.13294

106. Marshall NT, Stepien CA. Invasion genetics from thousands of larvae and eDNA of zebra and quagga mussels using targeted metabarcode high-throughput sequencing. Ecol Evol. In review.

107. Bubac CM, Spellman GM. How connectivity shapes genetic structure during range expansion; insights from the Virginia’s warbler. The Auk: Ornithological Advances. 2016; 133:213–230. https://doi.org/10.1642/AUK-15-124.1

108. Cristescu MEA, Witt JDS, Grigorovich A, Hebert PDN, MacIsaac HJ. Dispersal of the Ponto-Caspian amphipod Echinogammarus ischnus: invasion waves from the Pleistocene to the present. Heredity. 2004; 92:197–203. https://doi.org/10.1038/sj.hdy.6800395

109. Kolar CS, Chapman DC, Courtenay Jr. WR, Housel CM, Williams JD, Jennings DP. Bigheaded carps: a biological synopsis and environmental risk assessment. Bethesda, Maryland: American Fisheries Society Special Publication 33; 2007.

110. Liu J. Hypophthalmichthyinae. In: Pan JH, Zhong Y, Zheng CY, Wu HL, Liu JH (eds). The Freshwater Fishes of Guangdong Province. Guangzhou, China: Guangdong Science and Technology Press: 1991. 234–239.

111. Jerde CL, Chadderton WL, Mahon AR, Renshaw MA, Corush J, Budny ML, et al. Detection of Asian carp DNA as part of a Great Lakes basin-wide surveillance program. Can J Fish Aquat Sci. 2013; 70:1–5. https://doi.org/10.1139/cjfas-2012-0478

